# Conformational dynamics of Light-Harvesting Complex II in a native membrane environment

**DOI:** 10.1101/288860

**Authors:** F. Azadi-Chegeni, M.E. Ward, G. Perin, D. Simionato, T. Morosinotto, M. Baldus, A. Pandit

## Abstract

Photosynthetic light-harvesting complexes of higher plants, moss and green algae can undergo dynamic conformational transitions, which have been correlated to their ability to adapt to fluctuations in the light environment. Herein, we demonstrate the application of solid-state NMR spectroscopy on native, heterogeneous thylakoid membranes of *Chlamydomonas reinhardtii* (*Cr)* and on *Cr* Light-Harvesting Complex II (LHCII) in thylakoid lipid bilayers to detect LHCII conformational dynamics in its native membrane environment. We show that membrane-reconstituted LHCII contains selective sites that undergo fast, large-amplitude motions, including the phytol tails of two chlorophylls. Protein plasticity is also observed in the N-terminal stromal loop and in protein fragments facing the lumen, involving sites that stabilize the xanthophyll-cycle carotenoid violaxanthin and the two luteins. The results report on the intrinsic flexibility of LHCII pigment-protein complexes in a membrane environment, revealing putative sites for conformational switching. In thylakoid membranes, fast dynamics of protein and pigment sites is significantly reduced, which suggests that in their native organelle membranes, LHCII complexes are locked in specific conformational states.

**STATEMENT OF SIGNIFICANCE:** Photosynthetic Light-Harvesting Complexes undergo dynamic conformational transitions that regulate the capacity of the light-harvesting antenna. We demonstrate the application of solid-state (ss)NMR spectroscopy to investigate the structural dynamics of LHCII, the most abundant LHC complex of plants and algae, in native membranes. Selective dynamic protein and pigment residues are identified that are putative sites for a conformational switch.

## INTRODUCTION

Multi-pigment protein complexes in plants, moss, and photosynthetic algae perform delicate photo-physical and chemical tasks. To maintain a homeostatic balance in fluctuating sunlight conditions, these processes are tightly regulated by dynamic membrane responses to prevent photodamage (1, 2). The most abundant light-harvesting complexes, the peripheral antenna complexes LHCII, have been studied extensively for their molecular role in light-harvesting regulation. Through Raman spectroscopy of the LHCII xanthophylls it was revealed that isolated LHCIIs can undergo conformational changes *in vitro* and that they respond to excess light by undergoing similar conformational changes inside chloroplast organelle membranes, called thylakoids (3). Fluorescence studies have shown that individual LHCIIs have the capacity to switch between light-harvesting and photoprotective, excitation-quenched states (2-12). This concept has led to fluctuating-antenna models for describing excitation energy migration over a 2D protein lattice (13, 14). The LHCII crystal structures show selective sites with high b-factors. The structural dynamics of LHCII has been characterized with use of electron paramagnetic resonance (EPR) spin labeling combined with molecular modeling (MD) (4, 10, 15, 16). An MD study on monomeric LHCII in lipid membrane was also applied to correlate protein fluctuations to functional changes in pigment-pigment interactions (17). Those studies, however, do not capture the structural elements of native chloroplast membranes. In thylakoid membranes, many LHCIIs are held in specific arrangements within LHCII-Photosystem II super-complexes (18). The surrounding lipids and free xanthophylls that are present in thylakoid membranes further control the conformational dynamics of the membrane-embedded proteins (19). Furthermore, in living plants or algae reversible phosphorylation takes place during state transitions that involve membrane re-arrangements and a redistribution of LHCII over Photosystem I and Photosystem II (20).

The LHCII trimer complexes are isomers consisting of different pigment-bound polypeptides with overlapping sequences. The different polypeptides have been found to have distinct roles in photoprotection and state transitions (21, 22), suggesting that molecular recognition plays a role in LHCII regulatory functions. Phosphorylation will change the local structure of LHCII and may modify specific recognition sites (23). The phosphorylation sites are located in the N terminus of LHCII, which is not resolved in the crystal structures (10, 15). Protein-associated galactosyl lipids further have shown to play a role in stabilizing LHCII complexes and their interactions with partner proteins (24) as well as in controlling the supramolecular membrane architectures (25). Little is known about the functional roles of selective thylakoid lipids and how their dynamic interactions may contribute to the regulation of light harvesting.

Until now, no structure-based methods have been presented to experimentally probe the dynamics of LHCII inside a native lipid membrane environment. We here apply solid-state NMR (Nuclear Magnetic Resonance) spectroscopy to study the conformational dynamics of LHCII in lipid bilayers and in native thylakoid membranes. Solid-state NMR is a powerful tool for atomistic detection of membrane proteins and in some studies, NMR signals of membrane proteins could even be detected in native membrane or cellular environments, by using overexpression of the target proteins in prokaryotic and eukaryotic host-expression systems (26-32). In the following, we recorded NMR spectra of isolated LHCIIs that were reconstituted in lipid bilayers prepared from native-like lipid mixtures. In addition, we made use of the fact that LHCIIs are abundant in natural chloroplast membranes and performed NMR experiments directly on chloroplast thylakoid membranes containing LHCII that were isolated from wild type *U*-^13^C *Chlamydomonas reinhardtii* (*Cr*) cells. Strong overlap is observed between the spectrum of LHCII proteoliposomes and that of the thylakoid membranes, which allowed us to assess the dynamics of LHCII in its native environment.

## MATERIALS AND METHODS

### *Biosynthetic isotope labeling of* Cr *cells and sample preparation*

For the experiments on isolated Light-Harvesting Complex II (LHCII) trimers, *Cr* cells from strain *cw15* were cultivated in Erlenmeyer flasks with liquid tris-acetate phosphate (TAP) medium, at 100 rpm agitation and 21 °C in a growth chamber. For optimal labeling of the LHCII complexes, cells were grown under mixotrophic conditions under dim light to minimize the uptake of ^12^CO_2_ from air. Continuous illumination was provided from cool-white fluorescent lamps under low (< 25 μmoles photons m^-2^ s^-1^) photosynthetically active radiation (400-700 nm). The TAP medium (33) used to grow labeled cells, was prepared using ^13^C labeled sodium acetate and ^15^N labeled ammonium chloride (Sigma-Aldrich, Zwijndrecht, The Netherlands). Cultures in labeled medium were set up starting from an optical density at 750 nm (OD_750_) equal to 0.1 and were grown until OD_750_ = 1. Three rounds of cultivation in labeled medium were performed to ensure more than 95% labeling of the cells with ^13^C and ^15^N atoms. Thylakoid membranes were then isolated under previously described conditions (34). After the isolation, *Cr* thylakoids were re-suspended in buffer (50 mM Hepes-KOH pH 7.5, 5 mM MgCl_2_ with 50% glycerol). For isolation of the LHCII fractions, thylakoid membranes corresponding to 3mg/ml of total chlorophylls, according to the optical density at 680 nm, were washed with 50 mM ethylene diamine tetra-acetic acid (EDTA) and solubilized for 20 minutes on ice in 3 ml of final 1.2% n-Dodecyl α-D-maltoside ¢[-DM) in 10 mM Hepes (pH 7.5), after vortexing for 1 minute. The solubilized samples were centrifuged at 15000 x g for 30 minutes to eliminate any insolubilized material and the supernatant with the photosynthetic complexes was then fractionated by ultracentrifugation in a 0–1 M sucrose gradient containing 0.06% α-DM and 10 mM Hepes (pH 7.5), at 141000 x g for 40 hours at 4 □C. The green fraction (see Fig. 1D, SI section) corresponding to LHCII proteins was harvested with a syringe and Chl concentration adjusted to 2 mg/ml with buffer (50 mM Hepes, 5 mM MgCl_2_, pH 7.5). LHCII proteins solubilized in α-DM were reconstituted in lipid membranes whose composition mimics the native thylakoid membrane (47% monogalactosyldiacylglycerol (MGDG), 12% sulfoquinovosyldiacylglycerol (SQDG), 14% phosphatidylglycerol (PG) and 27% digalactosyldiacylglycerol (DGDG)) with a protein-to-lipid molar ratio of 1:55, according to the method described in Crisafi and Pandit (35). The chosen protein to lipid ratio is in the range of native protein packing densities in thylakoid membranes, where 70-80% of the membrane surface area is occupied with proteins (36).

**Figure 1.**
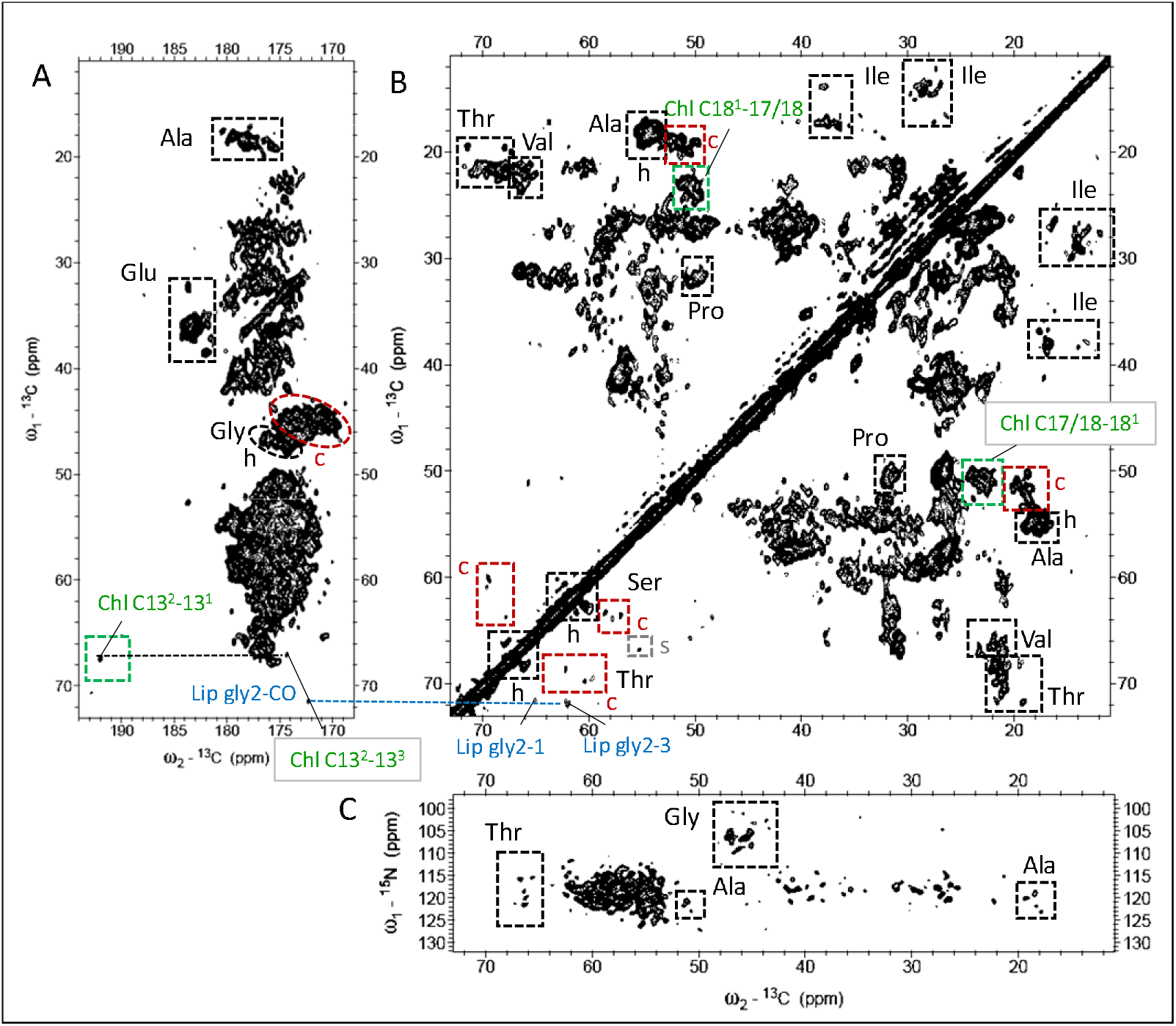
NMR spectra of LHCII proteoliposomes. A and B: ^13^C-^13^C CP-PARIS spectrum in the aliphatic (A) and carbonyl (B) region; C: ^15^N-^13^C NCACX spectrum. Identified clusters of amino-acid types are indicated with boxes. Helix (h), coil (c) and strand (s) correlations of Thr, Ser, Ala and Gly are indicated in black, red and gray in the ^13^C-^13^C spectrum. Chl and lipid (Lip) correlations are indicated in green and blue.

For experiments on whole fresh thylakoid membranes, *Cr* cells were cultivated in TAP medium using ^13^C labeled sodium acetate in a home-built photo chamber, under continuous illumination with cool white LEDs (~50 μmol m^-2^ s^-1^). Cells were harvested in the exponential growth phase, centrifuged and re-suspended in 0.2 volumes of MgCl2 buffer (1mM MgCl2, 0.1M HEPES, pH 7.5/KOH, 10% sucrose), and were ruptured by sonication on a 2500-Watt sonicator set at 10%. The isolation of fresh thylakoids was performed according to Chua and Bennoun (37) with some modifications. This procedure differed from the steps described above for LHCII isolation, by using sucrose gradient layers for purification of the thylakoid membranes in order to obtain more pure fractions. In the procedure, disrupted cells were overlaid with layers of sucrose (1 ml of 1.3 M, 1 ml of 0.5 M and 0 M sucrose). The gradients were ultra-centrifuged for one hour at 4 °C in a SW41 swing rotor (Beckmann) at 24 000 rpm (100000 g). The thylakoid fraction was isolated from the dark-green sucrose band (see Fig. S2A, SI section). With the used isolation procedure, membrane stacking is preserved.

For the LHCII NMR sample, 18 ml of LHCII in liposomes, containing approximately 10 mg LHCII and 1.5 mg Chl (as determined by OD_680_ of the Chls), was pelleted by ultracentrifugation (223 000 g, 4 °C, 90 min) and transferred to a thin-wall 3.2 mm solid-state NMR MAS (Magic Angle Spinning) rotor through centrifugation. For the thylakoid membrane NMR sample, 12 ml of freshly isolated thylakoid membrane containing 2 mg Chl and approximately 10 times more in protein content was pelleted by ultra-centrifugation (100 000 g, 4 °C, 45 min) and transferred to a thin-wall 3.2 mm MAS rotor.

### Gel electrophoresis

Coomassie-stained SDS-page was performed using 15% Tris-glycine gels (38). Samples were solubilized with a solubilization buffer (4 ×) containing 30% glycerol, 125 mM Tris pH 6.8, 0.1 M dithiothreitol, 9% SDS.

### Time-resolved and 77K fluorescence spectroscopy

Time-resolved fluorescence measurements on U-^13^C-^15^N LHCII in *α*-DM and on the LHCII proteoliposomes were performed using a FluoTime 200 (PicoQuant, Berlin Germany) time-correlated photon counter spectrometer. Samples were hold in a 1×1 cm quartz cuvette that was thermostated at 20 °C and excited at 440 nm using a diode laser (PicoQuant). 440 nm excitation. Fluorescence decay traces were fitted with multi-exponentials using a χ^2^ least-square fitting procedure. 77k fluorescence measurements were performed using a Fluoromax 3 spectrophotometer (Horiba, Jobin-Yvon, France). The samples were diluted in 50 mM 4-(2-hydroxyethyl)-1-piperazineethanesulfonic acid (HEPES), 5mM MgCl_2_ buffer and cooled in a nitrogen-bath cryostat to 77K. Excitation was performed at 440 nm and a bandwidth of 2 nm was used for selecting the excitation and emission wavelengths.

### Solid-state NMR experiments

Solid-state NMR spectra of *U*-^13^C-^15^N LHCII in proteoliposomes and of ^13^C-enriched thylakoid membranes were recorded on an ultra-high field 950-MHz ^1^H Larmor frequency spectrometer (Bruker, Biospin, Billerica, USA) equipped with a triple-channel (^1^H, ^13^C, ^15^N) 3.2 mm MAS probe. Typical π/2 pulses were 3 μs for ^1^H, 5 μs for ^13^C, and 8 μs for ^15^N. The ^1^H/^15^N and ^1^H/^13^C cross-polarization (CP) (39) contact times were 800 μs and 1 ms, respectively, with a constant radio frequency (rf) field of 35 and 50 kHz on nitrogen and carbon, respectively, while the proton lock field was ramped linearly around the n = 1 Hartmann/Hahn condition (40).The ^15^N/^13^Ca SPECIFIC-CP transfer (41) was implemented with an optimized contact time of 4.2 ms with a constant spin lock field of 2.5×ν_r_ applied on ^15^N, while the ^13^C field was ramped linearly (10% ramp) around 1.5×ν_r_. ^1^H decoupling during direct and indirect dimensions was performed using SPINAL64 (42) with ~83 kHz irradiation. The presented 2D ^13^C-^13^C PARIS (43) spectra were collected with a mixing time of 30 ms at 17 kHz MAS at a set temperature of −18 °C. The 2D NCA and NCACX experiments (44) were performed on the LHCII sample at 14 kHz MAS frequency and a readout temperature of −18 °C. For the NCACX experiment a PARIS (^13^C,^13^C) mixing time of 50 ms was used. The *J*-coupling based 2D ^13^C-^13^C INEPT-TOBSY (45, 46) experiments were recorded at −3 °C with a TOBSY mixing time of 6 ms at 14 kHz MAS. All spectra were processed using Bruker TopSpin software version 3.2 (Bruker, Biospin) with linear prediction and Fourier transformed in fqc (forward quadrature complex) mode. Spectra were analyzed by Sparky version 3.114 (47) and MestReNova 11.0 (Mestrelab Research SL, Santiago de Compostela, Spain).

### NMR chemical shift prediction using a plant-LHCII homology model

Homology models of *Cr* LHCII were built using the SWISS model web server (48) based on the LHCII crystal structure of spinach and using the Lhcmb1 or Lhcbm2 sequence of *Cr* LHCII (49). The Lhcbm sequences and respective PDB models were used as input for SHIFTX2 (50) in order to predict the ^13^C and ^15^N chemical shifts. ^13^C-^13^C predicted correlation spectra for use in Sparky (47) were generated by FANDAS (51).

## RESULTS

A biochemical analysis of our thylakoid preparations (Fig. S1 and S2, SI section) shows that the LHCII trimers are the most abundant pigment-containing complexes in the thylakoid membranes, while the higher-weight bands in the SDS page contain the photosystems, ATP synthase and cytochrome b_6_f complexes. Different molecular-weight bands are distinguished for LHCII due to the fact that the LHCII trimers are isomers consisting of different polypeptides of which the most abundant types are Lhcbm1, Lhcbm2/7 (Lhcbm2 and Lhcbm7 have identical mature peptide sequences) and Lhcbm3 (52). Our LHCII proteoliposomes contain the lipids MGDG (monogalactosyldiacylglycerol), DGDG (digalactosyl diacylglycerol), SQDG (sulfoquinovosyl diacylglycerol) and the phospholipid PG (phosphatidyl glycerol) at a protein to lipid ratio of 1:55 (*mol/mol*) to mimic the protein density and lipid composition of the native thylakoid membranes (53). The 77K fluorescence spectrum of the proteoliposomes shows a prominent band at 700 nm (Fig. S3, SI section), which is characteristic for LHCII aggregates. Aggregation of LHCII in liposome membranes is associated with the formation of quenched states (35, 54). Indeed, a fluorescence lifetime analysis gives an average lifetime of 0.7 ns for LHCII proteoliposomes compared to a lifetime of 3.5 ns for LHCII in α-DM detergent (Table S1 and Fig. S4, SI section).

Two complementary types of solid-state NMR experiments were employed to distinguish rigid and highly dynamic molecular species and protein subdomains. In Cross-Polarization (CP) based experiments, ^1^H-^13^C or ^13^C-^15^N magnetization is transferred via the dipolar interaction. Dipolar-based transfer experiments become inefficient for dynamic molecules or molecular segments that display sub-microsecond motions, as the dipolar couplings are averaged due to fast overall or local motion. Such dynamic species remain visible in NMR experiments such as INEPT (10) and -TOBSY (9) where magnetization is transferred via *J* couplings. The combination of such *J*-based pulse sequences can hence be employed for screening very flexible parts of large biomolecules that undergo large-amplitude motions on ps-μs time scales (55). On the other hand, membrane proteins typically contain long transmembrane domains with limited flexibility, which are predominantly detected in dipolar-based experiments (55).

Dipolar-based ^13^C-^13^C (CC) and ^15^N-^13^C (NC) experiments of LHCII in proteoliposomes are shown in Fig. 1 for the protein regions the selective pigment and lipid regions are shown and discussed further down in Fig. 3. The CP-PARIS CC spectrum verifies that our sample contains pigment-protein complexes of which all components are uniformly labeled. For clarity, the full spectrum in the aliphatic region without annotations is also shown in Fig. S5 in the SI section. Resonance signals of specific amino-acids, i.e. the Cα-Cβ correlations of threonine (Thr), serine (Ser), alanine (Ala) and the Cα-C’ correlations of glycine (Gly), can be identified and classified as helix or coil based on their unique chemical-shift patterns and clear spectral separation of their Cα-Cβ (Thr, Ser, Ala) or Cα-C’ (Gly) correlation signals in the CC spectrum. We use those amino acids as spectroscopic markers. To evaluate if their correlation patterns agree with the known secondary structure of LHCII, integrated NMR signal intensities in helix and coil regions of Thr, Ser, Ala and Gly are presented in Fig. 2. The NMR-estimated coil and helix contributions are compared to the predicted helix and coil contents according to the amino acid sequence and according to a structural model that was build based on the plant LHCII crystal structures (10, 15) using the amino-acid sequence of Lhcbm1, one the most abundant polypeptides of *Cr* LHCII.

**Figure 2.**
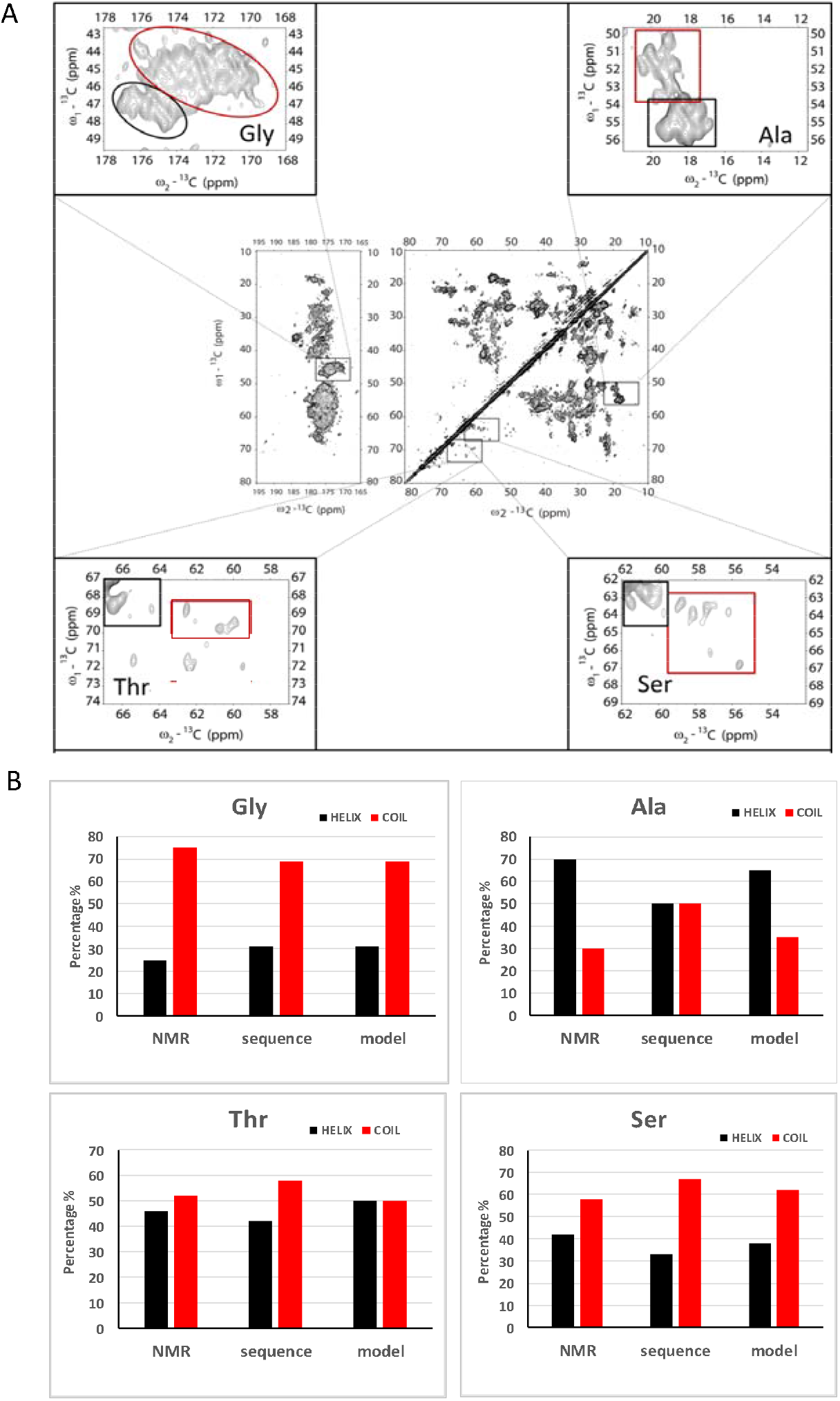
NMR-estimated helix and coil percentages of Gly, Ala, Thr and Ser residues. A: Insets of the ^13^C-^13^C PARIS spectrum containing Gly, Ala, Thr or Ser correlation signals. Helix and coil regions are indicated with black and red boxes respectively. B: Helix and coil percentages according to the NMR, Lhcbm1 sequence and Cr LHCII homology model.

**Figure 3.**
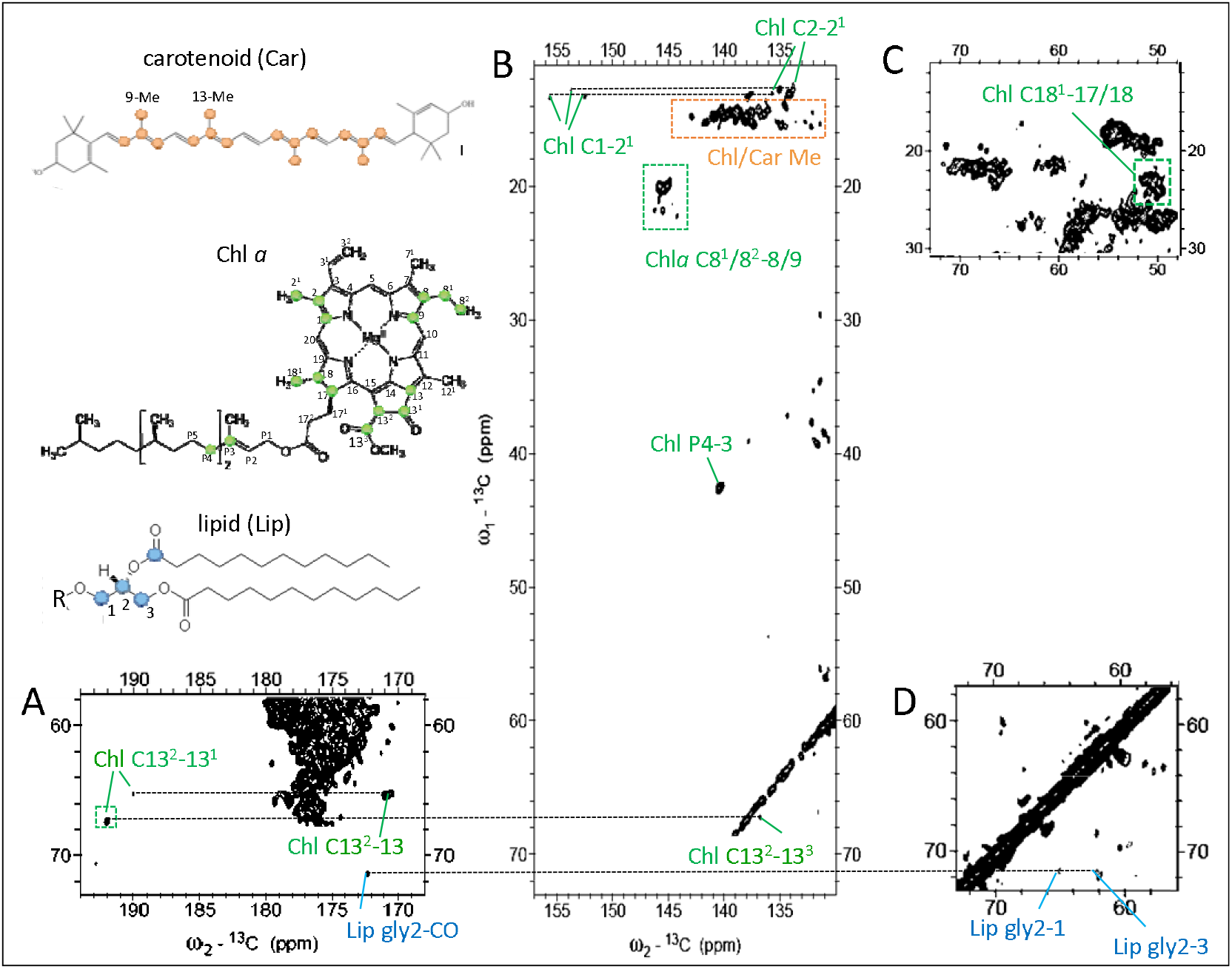
Pigment and lipid correlations in the PARIS ^13^C-^13^C NMR spectrum of LHCII proteoliposomes. Insets of the carbonyl (A), aromatic (B) and aliphatic (C and D) region with identified chlorophyll (Chl), carotenoid (Car) and lipid (Lip) correlations indicated in green, orange and blue. The structures of carotenoid (lutein), of Chl a and of a glyco-or phospholipid are drawn in the left side of the figure with their attributed carbon atom types color-coded.

The structural model lacks the first 13 residues in the N terminus of LHCII as this part is not resolved in the LHCII crystal structures and therefore has lower numbers of coil amino acids than the full Lhcbm1 sequence. Comparing the NMR-estimated helix and coil contents of Gly, which amino acid type are abundant in LHCII and evenly distributed over the Lhcbm1 sequence (See Fig. S6, SI section), the estimated fractions agree with those anticipated and confirm that the LHCII complexes have well-folded secondary structures. For Ser and Thr, the NMR-estimated contents also reasonably agree with the anticipated numbers. One Thr is at the edge of a 310 helix fragment on the luminal site and may adopt a coil conformation. For Ala, the NMR-estimated helix and coil contents agree with the model but the estimated coil content is lower than anticipated based on the full sequence. The full sequence contains additional Ser, Thr and Ala amino acids in the N terminus (2 x S, 2 x T and 3 x A). We suspect that the flexible N tail is not detected in the NMR CP-PARIS spectrum, in agreement with earlier work on ^13^C-^15^N-Arg *Cr* LHCII, where the number of detected Arg signals matches with the anticipated number based on the structure if the N terminal part was left out (56). No Gly amino acids are found in the N terminus of Lhcbm1 so that the model and the full sequence contain the same helix and coil Gly contents. In Fig. S7 in the SI section, NMR-estimated helix and coil contents are also compared to those calculated from the structure of Lhcbm2/7 and to the model and full sequence of Lhcbm3. Lower agreement is observed for the Ala and Ser helix/coil contents, suggesting that in our LHCII preparations Lhcbm1 is dominant.

In addition to signals originating from the protein content, Chl and carotenoid signals can be identified in regions of the CC spectrum where no protein signals are expected. They are shown in Fig. 3, which shows enlarged regions of the LHCII CC spectrum where Chl, carotenoid and lipid signals occur. Signals are identified from several Chl macrocycle ring carbons (C1, C2, C8, C9, C13, C13^1^, C13^2^, C17/18) and their neighboring side-chain atoms (C2^1^, C8^1^, C8^2^, C13^3^, C18^1^). The C8, C9, C8^1^, C8^2^ signals are specific for the side chain Chl *a*, while the other signals could be either from Chl *a* or *b*. In addition, signals are detected of atoms P3 and P4 in the Chl tail. Strong signals of carotenoids (Car) are observed that are easily distinguished from lipid signals owing to their correlations between the conjugated chain and Me resonances of carotenoids, accumulating between 15-19 ppm (ω1) and 130-145 ppm (ω2) (Fig. 3B). The Chl correlations of macrocycle ring atoms with side chain atoms C7^1^ and 12^1^ also fall in this region. A single set of lipid glycerol carbon resonances is observed. It is well-known that each monomer unit of LHCII contains one phosphatidyl-glycerol (PG) lipid, which forms the ligand for Chl611 (nomenclature Liu et al. (15)). Based on their chemical shifts, we attribute the detected lipid resonances to the head carbons of LHCII-bound PG lipid molecules. From its appearance in a CP-based spectrum it can be deduced that this lipid does not exchange on a typical NMR time scale (~0.1 ms) and is a structural lipid.

We now compare the NMR spectrum of LHCII in liposomes with the NMR spectrum of thylakoid membranes containing LHCII. To assess the overall dynamics of our proteoliposome and thylakoid membrane sample preparations, we first compared the relative intensities of 1D ^13^C MAS-NMR spectra obtained through ^1^H-^13^C cross-polarization (CP) to those obtained through a 90° pulse with direct polarization (DP) of the carbons (Fig. S8, SI section). The spectra were recorded at −3 °C to include NMR signals of mobile sites. For pure solids, the CP signal intensities will be ~4 times enhanced compared to the DP signal intensities, based on the gyromagnetic ratio of ^1^H relative to ^13^C whereas for dynamic molecules the relative CP signal intensities will be lower as polarization transfer becomes less efficient. For both samples, the CP and DP signal intensities in the aliphatic region (0-60 ppm) that include the Cα, Cβ and other amino acid side chain atom signals, are similar, while the DP spectrum is enhanced in the aromatic (110-130 ppm) and carbonyl (170-180 ppm) region. The additional peaks in the DP spectrum contain those of fatty acids and carbohydrate atoms from the galactosyl lipid heads. The CP/DP ratios reflect the state of matter of the sample preparations and indicate that the samples are not pure solids (which would have given ~4 times signal enhancement for CP) but form gel-like states as expected. From their similar CP/DP relative intensities, we conclude that the overall dynamics of the two samples are comparable.

Fig. 4 shows the 2D ^13^C-^13^C CP-PARIS spectrum of LHCII (black) overlaid with the spectrum of thylakoid membranes (red) that was recorded and processed under identical conditions. Strong overlap of the two spectra demonstrates, in agreement with the biochemical analysis, that the most abundant signals that dominate the 2D ^13^C-^13^C thylakoid spectrum arise from LHCII. Contributions of other membrane components are visible as additional signals in the thylakoid spectrum compared to the overlaid LHCII spectrum. Strong resonance correlations accumulate between 70-80 ppm that can be assigned to the galactosyl heads of thylakoid lipids. Fig. 5 show the overlaid spectra of LHCII in liposomes (black) and thylakoid membranes (red) focusing on the Ala (A), Ser (B) and Thr (C) selective spectral regions. Fig. S9, SI section shows the spectra in the aromatic region where Chl and carotenoid signals appear. Note that the pigment compositions and stoichiometric contents in the thylakoid membranes differ from those of isolated LHCII because the thylakoid contains additional photo-complexes with different pigment types and stoichiometries. The LHCII relative protein content in thylakoid membranes therefore differs from the LHCII relative pigment contents.

**Figure 4.**
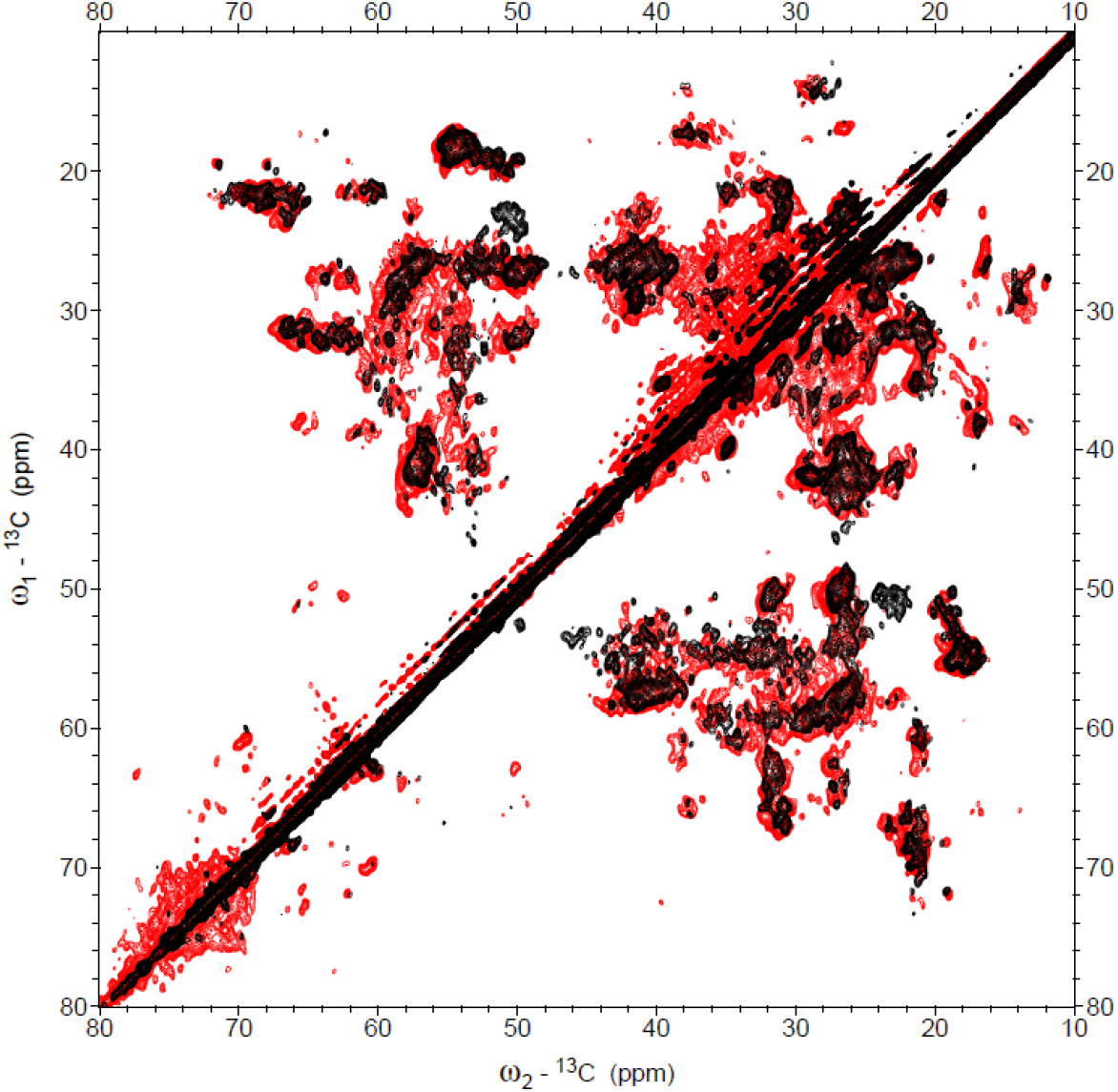
NMR comparison of LHCII proteoliposomes and thylakoid membranes. ^13^C-^13^C CP-PARIS spectrum of thylakoid membranes (red) with the LHCII spectrum (black) overlaid.

**Figure 5.**
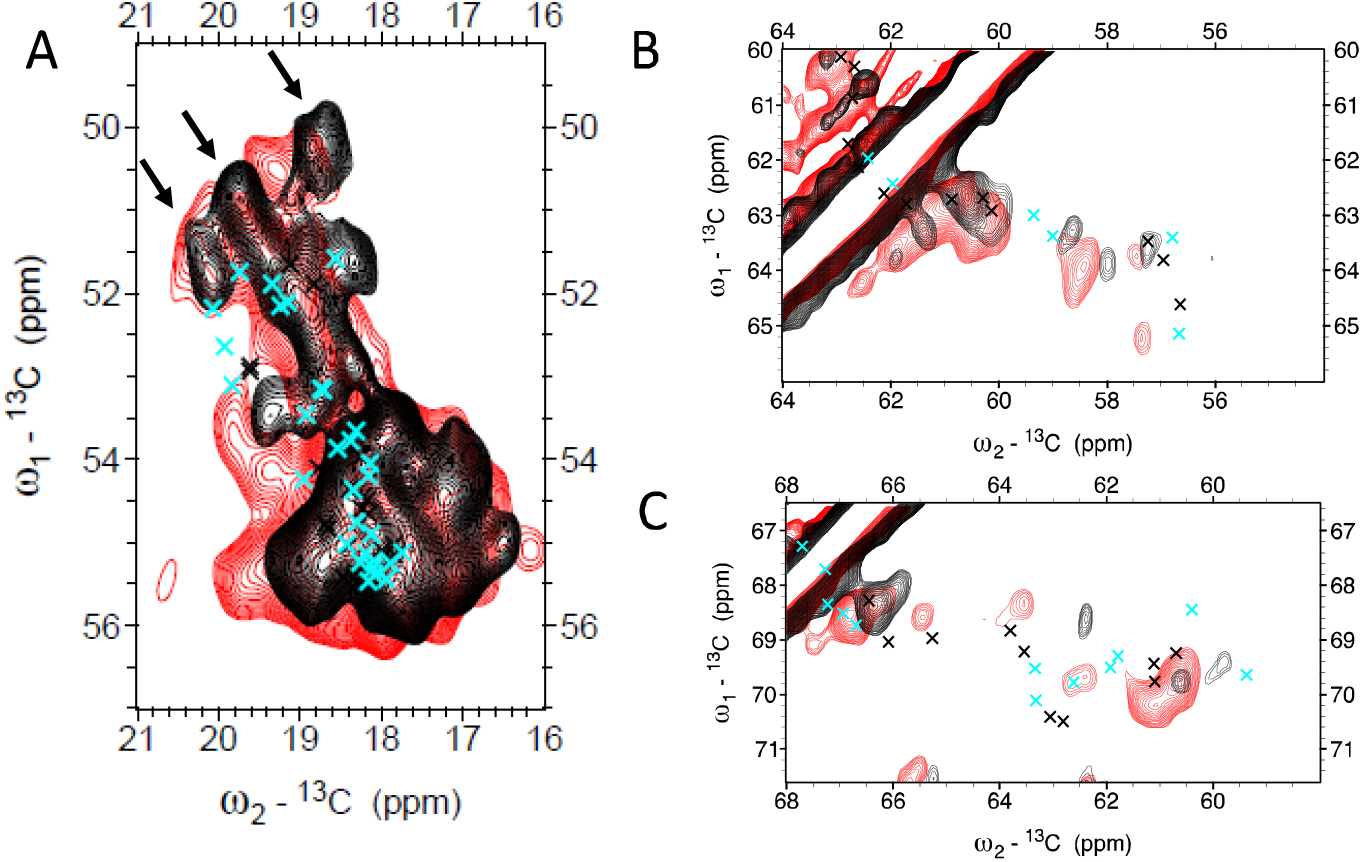
^13^C-^13^C CP-PARIS spectrum of thylakoid membranes (red) with the LHCII spectrum (black) overlaid in the Ala (A), Ser (B) and Thr (C) spectral regions. Chemical-shift predictions of Lhcbm1 (black crosses) and Lhcbm2/7 (cyan crosses) are overlaid.

Representative 1D slices of the spectral regions are also shown Fig. S9, SI section to indicate the signal to noise levels at different ω1 frequencies. While most of the Ala signals in the two spectra overlap, low agreement is seen comparing their Ser and Thr coil signals. The spectra in Fig. 5 are overlaid with predicted Cα and Cβ chemical-shift correlations that were generated from the Lhcbm1 and Lhcbm2/7 homology models using the program SHIFTX2 (50) and simulated as CC spectrum using the program FANDAS (51).

The full CC spectrum of LHCII with all the predictions overlaid can be found in Fig. S10 and an NC spectrum with overlaid predictions can be found in Fig. S11, SI section. Overall the pattern of experimental correlations matches well with the predictions. However, in the selective regions we observe several deviations. Remarkably, three strong Ala peaks appear in the coil region of the spectrum in Fig. 5A (indicated by the arrows for the LHCII spectrum) that are better resolved in the proteoliposome spectrum and of which the Ala Cα frequencies are shifted upfield compared to the predicted peaks. In the Thr region (Fig. 5C) predicted peaks aare close to experimental Cα-Cβ cross-correlation signals in the thylakoid spectrum, but are not close to any experimental peak in the LHCII proteoliposome spectrum. None of the evaluated residues are in Van der Waals contact with pigment ligands, excluding direct contact interactions as the cause of chemical-shift anomalies. In the pigment region, we note that the signals that we attributed to Chl C18^1^-C17 or C18^1^-18 correlations in the aliphatic region of the CC spectrum of LHCII proteoliposomes (see Fig. 3) are not visible in the spectrum of thylakoid membranes (Fig. S9, SI section). In contrast, Chl signals of ring carbons resonating in the region 130-145 ppm that correlate with the side chain Me signals in the region 10-20 ppm appear in both spectra.

To explore the presence of dynamic protein sites, we proceeded with *J*-based (INEPT-TOBSY) experiments that are exclusively selective for molecules with strong dynamics and large-amplitude motions. To enhance the mobility and emphasize signals from very flexible regions, the INEPT-TOBSY experiments were carried out at were recorded at a higher readout temperature of −3 °C. Fig. 6 presents the 2D INEPT-TOBSY spectrum of the proteoliposomes (black) overlaid on the INEPT-TOBSY spectrum of thylakoid membranes (red). As expected by the fact that the majority of the protein correlations are detected in the dipolar-based spectra, only a limited set of protein signals are detected that we refer to as *“J”* amino acids. The overlaid spectra clearly show that many more *J* signals are detected in the INEPT-TOBSY spectrum of LHCII (black) than in the spectrum of thylakoid membranes (red).

**Figure 6.**
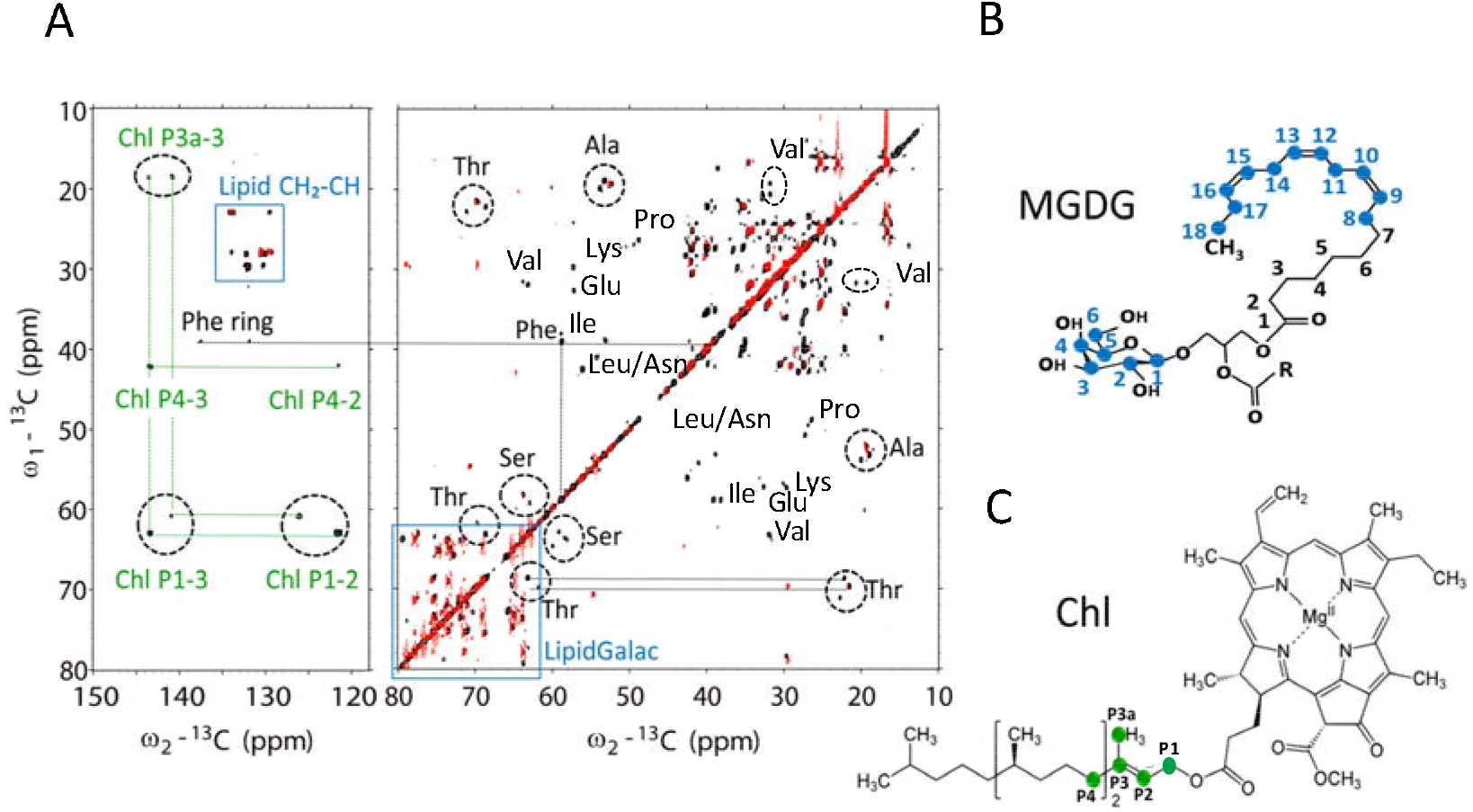
J-analysis of LHCII proteoliposomes and thylakoid membranes. A: Overlay of the INEPT-TOBSY spectrum of LHCII (black) and of thylakoid membranes (red). Resonance signals of Chl phytol chains, lipids and Ala, Thr, Ser, and Phe amino acids are indicated. B: chemical structure of MGDG with assigned carbon atom types colored in blue. C: chemical structure of Chl with the assigned carbon atom types colored in green.

We could assign the *J* residues in the LHCII spectrum to Ala, Thr, Ser, Phe, Pro, Val, Lys, Ile, Glu, Asn and Leu amino acid types (see Table S2 and Fig. S11, SI section for the chemical-shift assignments). Based on the Cα and Cβ chemical shifts, the *J* amino acids can all be classified as having non-helical structure. Because they represent amino acids that display fast and large-amplitude motions, we expect them to be located in the non-helical stretches and in the N- and C-termini. Remarkably, whereas random coil correlations often overlap, here distinct chemical shift peaks are observed for *J* signals of same amino acid types. For instance, we see three distinct Ala Cα-Cβ correlation peaks in the spectrum. This indicates that the *J* amino acid containing segments are dynamical, but form structured elements. The INEPT-TOBSY spectrum of LHCII also contains two sets of Chl phytol chain signals (chemical shift assignments in Table S3), revealing that two Chls have dynamic tails. In the thylakoid INEPT-TOBSY spectrum, no Chl correlations are observed.

The lipid signals in the NMR spectra of LHCII allow us to identify the nature of the LHCII-associated lipid molecules. The ^13^C lipid NMR signals must originate from the original thylakoid lipids that remained associated with the LHCII complex after purification, because the reconstituted lipids are not isotope labeled. Several galactolipids are detected in the INEPT TOBSY spectrum that can be assigned to MGDG or DGDG (57). Their chemical shift assignments are presented in Table S4 and the connectivities are drawn in Fig. S12 in the SI section.

The lipid signals in the thylakoid spectra provide a unique molecular picture of lipid dynamics inside the thylakoid membrane. Lipid signals are detected both in the *J*-based and in the dipolar-based CC spectrum, indicating that the thylakoid membrane contains mobile lipids with strong dynamics and low segmental order, as well as immobilized lipids with low dynamics, which notion is in agreement with our previous work (58). In the previous work, we tentatively assigned lipid signals from 1D ^13^C MAS NMR (59). Owing to the higher resolution and 2D spectra in our present study, we now can distinguish different lipid types via differences in the sugar head groups of MGDG, DGDG and SQDG and make a connecting walk through the ^13^C-^13^C spectra to correlate CH-CH2 correlations. The thylakoid membrane contains lipids with high degree of unsaturation. Their CH-CH_2_ correlations involving double-bonded carbons exclusively appear in the INEPT spectrum and not in the CP-based spectrum, revealing that the unsaturated lipids in the thylakoid membrane have highly mobile tails.

## DISCUSSION

### Dynamic sites in LHCII according to J-based NMR spectroscopy

The majority of NMR signals of LHCII proteoliposomes appear in the CP-based spectrum whereas only a selective set of signals is seen in the *J*-based spectrum, verifying that the LHCII complexes are stabilized in the lipid membranes. The *J*-base INEPT-TOBSY spectra reveal selective rapid, large-amplitude motions on a ps-ns time scale, revealing intrinsic strong dynamics of these protein and pigment sites. According to the 77K fluorescence spectrum and fluorescence lifetime analysis, the LHCIIs form aggregate states inside the liposome membranes. We conclude that despite their strong aggregation, selective sites in liposome-reconstituted LHCIIs have considerable dynamics. The INEPT-TOBSY spectrum was collected at −3 °C while the CP spectrum was collected at −18 °C. Because rigid amino acids that are detected in the CP-based spectrum at −18 °C may become dynamic when the temperature is raised to −3 °C, the CP-PARIS and INEPT-TOBSY spectrum of LHCII are overlaid in Fig. S13 in the SI section. One Ser *J* signal indeed overlaps with a correlation in the CP-based spectrum and one Ala *J* signal overlaps, indicating that those two amino acids are rigid at −18 °C but are dynamic at −3 °C. Other *J* signals in the INEPT-TOBSY spectrum do not overlap with any of the protein signals seen in the CP-based spectrum and must arise from different amino acid residues. Those could be amino acids in the N terminus as we suspect, supported by analysis of the integrated the helix and coil intensities and by earlier work (56) that amino acids in the N terminus are not detected in the CP-based spectrum. The N terminal domain part of Lhcbm1 is VEARRTVKPASKASTPD, which matches with many of the observed *J* residue types (3 x A, 3 x S, 3 x P, 3 x T, 1 or 2 L and 1 x V, I, F, K, E and N amino acid signals). For Lhcbm3 the equivalent stretch is KATGKKGTGKTAAKQAPASSG while the sequence of Lhcbm2 the N-terminal domain is only IA. The N terminus could not be resolved in plant LHCII crystal structures, which was ascribed to its non-uniform positioning in the crystals (10, 15). To resolve the structure of this protein stretch, Fehr et al. modeled the N-terminal section based on distance mapping using EPR spin labels. They showed that the N-terminal domain is dynamic, but does not have a random structure and covers only a restricted area above the superhelix in LHCII (16). Our notion that *J* amino acids form highly dynamical, but structured, elements agrees with their model. In a coarsegrained simulation of LHCII in thylakoid lipid bilayer, the N terminus was shown to be preferentially located on top of the membrane-protein interface, enabling interaction with the lipid headgroups (60).

In addition to the N-terminal loop, the C terminus is predicted to be very dynamic for lipid-embedded LHCII trimers according to coarse-grain MD simulations (60). The flexible part of the C terminus contains the stretch TKFTPQ (Lhcbm1) or TKFTPSA (Lhcbm2) involving the two Thr amino acids T213 and T216. One of the observed *J* amino acids is a phenylalanine (Phe), while there are no Phe amino acids in the N terminus. Therefore, some of the *J* amino acids likely belong to the C terminus.

Inspection of the LHCII crystal structures shows that Chl 605 and 606 (nomenclature according to Liu et al. (15)) are the only Chls in the structure with unresolved phytol chains, emphasizing their dynamics, whereas the other phytol chains are resolved and form tight interactions within the pigment-protein complex. The chains of Chl 605 and 606 are oriented outward according to the orientation and positioning of their macrocycle rings in the structure and likely will not be motion-limited by intra-complex interactions. In fact, Chl 606 is ligated by a water molecule and could be very dynamic. In a coarse-grain model of LHCII in thylakoid lipid bilayer, the phytol tail ends of Chl605 and 606 undergo the largest fluctuations (60). We attribute the two sets of Chl phytol resonances in the INEPT TOBSY spectrum to Chl 606 and 605. The CP-PARIS spectrum shows correlations between P3 and P4, which are the two phytol atoms close to the Chl ring. No other phytol chain atom signals are observed in the CP-based spectrum. In previous studies we found that the dynamics of protein-bound chromophores often fall in an intermediate regime where their 13 carbon signals are neither detected in CP-, nor in INEPT-based NMR experiments (8). Indeed, new cross correlation peaks of carbons along the Chl phytol chain are seen in a CC spectrum that is obtained by direct excitation (DP) as illustrated in Fig. S14 in the SI section.

The INEPT-TOBSY spectrum of LHCII proteoliposomes further reveals that the galactosyl membrane lipids are associated with LHCII are very dynamic. Because of their isotope labels, they are intrinsic thylakoid lipids that must have been purified together with the LHCII complexes. We assume that they are co-purified annular lipids that in the original thylakoid membrane formed a shell around the LHCII complexes. In particular MGDG lipids have been shown to interact with the LHCII complexes (60, 61). Upon reconstitution of LHCII, those lipids may have exchanged with the liposome bulk lipids. In contrast, a PG lipid is observed in the CP-PARIS spectrum, consistent with the fact that this is a structural lipid that stabilizes the LHCII trimer conformation and does not exchange on NMR time scale. MD simulations of LHCII monomers embedded in a lipid bilayer performed by Liguori et al. predict that flexibility of the N terminal stretch induces significant disorder of the PG molecule that ligates Chl611 and that is stabilized by a conserved Tyr (Y31 in the Lhcbm1 sequence) (9). This is not in agreement with what is observed in our experiments. In the dipolar-based NMR spectrum the correlating set of PG lipid signals is well resolved (see Fig. 3) implying that this lipid has a steady conformation. The discrepancy may be explained by the fact that the MD simulations were performed on LHCII monomers, whereas our samples contained LHCII trimers. The PG structural lipid is located at the interface between the monomers and possibly forms a more rigid structure in trimeric LHCII.

### Flexible sites in LHCII according to dipolar-based NMR spectroscopy

Comparing the CP-PARIS spectrum of LHCII proteoliposomes to the CP-PARIS spectrum of thylakoid membranes containing LHCII, poor overlap is observed in the Thr and Ser spectral coil regions and the correlation peaks in the LHCII proteoliposome spectrum are weak, suggesting that Thr and Ser correlations in the coil regions are not well resolved. Two Thr amino acids, T213 and T216, are located at the C terminal stretch that is predicted to be highly dynamic. Yet, the integrated helix and coil Thr peak intensities of the LHCII proteoliposome spectrum match quite well with those anticipated. This discrepancy may be solved if we take a closer look at the Thr amino acids in the LHCII structure. We considered T188 and T205 as helical, whereas T205 is part of a 3_10_ helix and T188 is at the lumen edge of helix A. In fact, according to the Lhcbm1 generated NMR predictions, the NMR Cα and Cβ correlations of T188 and T205 fall into the coil region. If T188 and T205 are considered as coil, the anticipated fraction of Thr signals in the coil region is 70%, which would be significantly higher than estimated from the coil contributions in the experimental spectrum.

To identify NMR chemical-shift anomalies, we consider deviations between predicted (ω pred) and experimentally observed (ω exp) Cα-Cβ or NH-Cα protein backbone correlations significant if √[wω_1_ exp - ω_1_ pred)^2^ + (ω_2_ exp-ω_2_ pred)^2^] ≥ 1.2 ppm. (62). In the LHCII proteoliposome spectrum, anomalies in the Ser and Thr coil regions are observed for the residues T16, T35, T188, T205 and S110 (in Lhcbm1) and T22, T38, S113 and T191 (in Lhcbm2) as shown in Fig. S15. Additional anomalies are seen in the NCA spectrum for V106, I111 and T213 and are shown in Fig. S11. For the highlighted amino acids, backbone chemical shifts either fall into a different region than predicted or are lacking in the LHCII spectrum. In contrast, in the thylakoid spectrum, matching correlation signals are found. In other words, coil correlation signals of LHCII in thylakoid membranes match with structurebased predictions, while selective coil signals of LHCII in proteoliposomes are invisible or falling into different regions. Intermediate protein dynamics could cause invisibility of signals in CP-based spectra as thermal motions on a microsecond time scale lower the efficiency of cross polarization. In addition, dynamic solvent-exposed residues may have multiple conformers at low temperature when their motions are frozen out, resulting in inhomogeneous line broadening or invisibility (63). Fig. 7 presents the amino acids with anomalous shifts and their location in the Lhcbm1 or Lhcbm2/7 sequences and Lhcbm1 structure. Anomalies are found for residues in the stromal loop, the luminal edge of helix C and helix A and for residues close to the C terminus. Indeed, a CG protein model of LHCII trimers predict that the C terminus, stromal loop and edges of the AC and BC loops have the highest B-factors (60). In a previous study we found that an arginine in the stromal loop adopts a different structure in aggregated LHCII than in frozen solutions of LHCII trimers in detergent micelles (56). The notion of a flexible stromal loop is consistent with the observations in present work, where we find Thr chemical-shift anomalies in the same loop segment.

**Figure 7.**
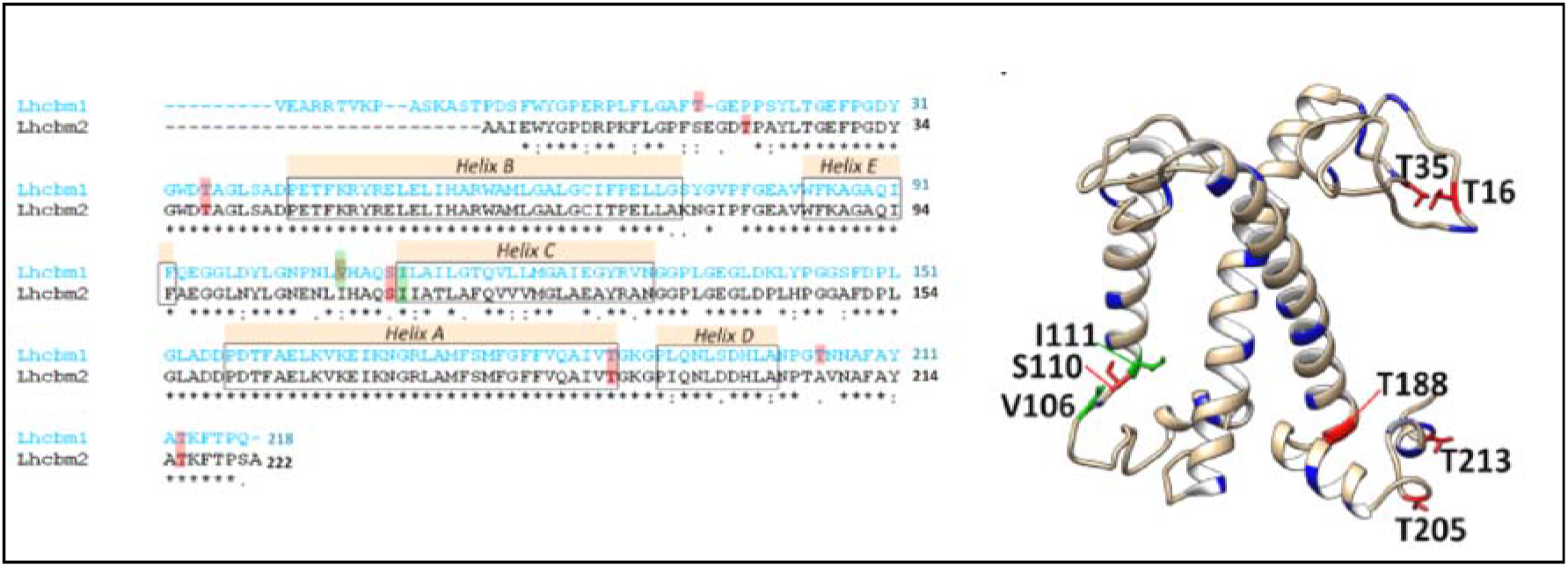
Protein sites with deviating NMR chemical shifts. Left: amino-acid sequences of Lhcbm1 and Lhcbm2 with deviating shift residues indicated. Right: homology model of Lhcbm1, highlighting Thr, Ser and Ala residues with matching chemical shifts in blue and residues with deviating NMR chemical shifts in red (deviations in CC spectrum) or green (deviations in NC spectrum).

Several of the amino acids with anomalous shifts or weak signals are close to pigment binding sites as illustrated in Fig. 8. We speculate that flexibility of the protein matrix at these spots could influence the pigment properties. It is known that the loop fragment containing T35 controls the position and orientation of L2 carotenoids and in the structure T35 is close to the head group of Lutein 2 (10, 15). The geometries of LHCII carotenoids in the lutein pocket are found to tune light-harvesting efficiency (64) and structural changes of those sites could permit access to different dark states (65). At the lumen site of LHCII, S110 and I111 control the position and dynamics of Chl605 that is ligated to I111 and of Chl606, which are the Chl molecules with fast moving tails. T205 connects the amphipatic helix D to a 3_10_ helix fragment at the C terminus of LHCII containing T213 that together with the stromal loop stretch around T16 stabilizes the xanthophyll-cycle carotenoid violaxanthin. Plasticity of the V1 pocket could be a requirement for LHCII to bind and release carotenoids under influence of (de) epoxidase enzymes during the xanthophyll cycle and explain why V1 carotenoids are loosely bound and easily removed during protein isolation.

**Figure 8.**
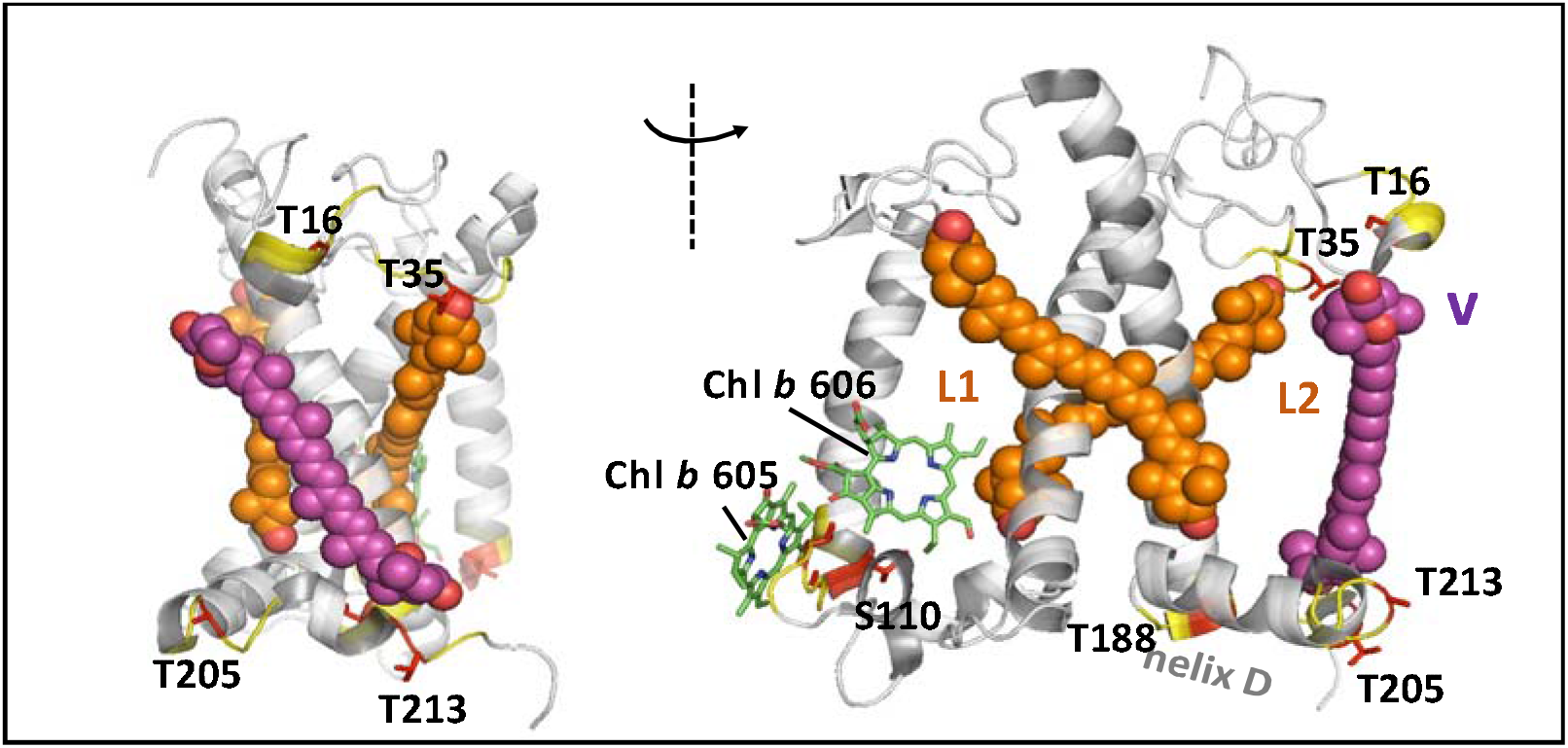
LHCII sites with deviating shifts and pigments in close proximity. Side view (left) and front view right) of spinach LHCII with Lutein 1 (L1), Lutein 2 (L2), violaxanthin (V) represented as spheres and Chl605 and 606 as sticks. Deviating residues are highlighted in red and adjacent residues (residue i±1, i±2) are colored yellow.

### Dynamics of LHCII in native thylakoid membranes

The NMR spectrum of thylakoid membranes containing LHCII provides us with a view on the dynamics of LHCII in a native membrane environment. We observe that fast, large-amplitude motions are suppressed for LHCII in native thylakoid membranes. The INEPT-TOBSY spectrum of the native thylakoid membranes lacks the protein *J* signals that are present in the LHCII proteoliposome spectrum and also lacks the signals of two mobile Chl tails. As strong overlapping signals are observed in dipolar-based correlation spectra of both samples, the absence of *J* signals in the thylakoid spectrum cannot be attributed to a lower LHCII content in the thylakoid sample. Comparable CP and DP spectral intensities further indicate that the overall phase of matter conditions of the two sample preparations are comparable and excludes a trivial explanation that the thylakoid sample overall is more rigidified.

The cross-correlation peaks that we attributed to Chl C17/18-C18^1^ correlations in the CP-based spectrum of LHCII proteoliposomes are not visible in the thylakoid spectrum. We presume that this has to do with differential dynamics of the Chls in proteoliposomes and in thylakoid membranes with respect to the 17, 18 and 18^1^ macrocycle atoms, which form the ring site that is connected to the phytol chain. In an earlier study we noted that the visibility of the LHCII Chl double-bonded ring atoms in CP-based NMR spectra were strongly dependent on temperature conditions and disappeared when the temperature was raised from −50 °C to −30 °C (8).

Taken together, a picture emerges for the intrinsic dynamics in LHCII. The N-terminal tail and C-terminus of LHCII can undergo selective, fast, segmental motions on (sub-) nanosecond timescales, while the connected stromal loop, the EC loop and luminal sites, involving sites that are close to carotenoid binding pockets, have conformational flexibility and may undergo thermal equilibrium motions on a microsecond timescale. In native thylakoid membranes, LHCII fast segmental motions are suppressed and flexible sites have differential dynamics or adapt a conformation that differs from liposome-embedded LHCII.

To explain the differences, we consider the supramolecular interactions that LHCII complexes can have inside the thylakoid membranes. Thylakoid membranes form stacks that are stabilized by transversal salt bridges between the stromal sites of LHCIIs (the N-terminal site) (34) that could significantly reduce the dynamics of the N terminus, compared to liposome-embedded LHCII where the N terminus will be exposed to the lipid-water interface. Phosphorylation of specific Thr in the N-terminus of LHCII may constrain dynamics of the N tail in thylakoid membranes and could modify the structure of the connected stromal loop.

Recent cryo-electron microscopy (cryo-EM) structures of *Cr* PSII-LHCII supercomplexes show extensive interactions between LHCIIs and the monomeric complexes CP43, CP47, CP26, CP29 and protein PsbW (66, 67). Strongly-bound LHCII (S-LHCII) in the super-complex interacts with CP43, CP26 and the protein PsbW via the AC and EC loops. Loosely bound LHCII (N-LHCII) interacts with CP47 and PsbW via Chl611 on the stromal side and Chl614 and short helix D on the luminal side. Moderately-bound LHCII (M-LHCII) and N-LHCII are tightly associated with CP29 at the stromal side via extensive interactions at the AC loop, Chl608 and neoxanthin and at the luminal side via Chl605 and the BC loop. Multiple interactions are observed among the LHCIIs via the amphipatic helix D that bring Chl605 and Chl606 of connected LHCIIs close together. The phytol chain of Chl605 that is lacking in LHCII crystal structures is partly resolved in the *Cr* LHCII-PSII structure. Interactions involving helix D could influence the conformation and dynamics of T188 and T205 at the lumen side while the interactions involving the BC loop are close to V106, S110 and I111. With our chemical-shift analysis, we don’t have suitable NMR reporters at the AC loop. The AC loop of Lhcbm1 contains one Ser (S147), one Ala (A153) and a number of Gly residues. Predicted Cα-Cβ correlations of Ser and Ala however fall in regions where multiple signals accumulate and also the Gly Cα-CO coil correlations are not dispersed. Whereas reduced dynamics of LHCII protein and pigment sites can be explained by their interactions in super-complexes, thylakoid membranes also contain a pool of free LHCIIs. The thylakoid INEPT-TOBSY spectrum, which has excellent signal to noise, does not show weak signals from a subset of free LHCIIs. This suggests that also for free complexes, fast protein and pigment fluctuations are suppressed by the surroundings of the thylakoid environment.

The thylakoid INEPT-based CC spectrum contains only few protein signals, but contains multiple signals of unsaturated thylakoidal lipids with highly mobile tails. This could point to a phase separation between dynamical lipid-rich and rigid protein-rich membrane regions. Lipid polymorphism has been revealed in chloroplast thylakoid membranes, including the reversible formation of inverted hexagonal phase structures by the non-bilayer lipid MGDG (68).

Finally, our findings call to question whether or not spontaneous fluctuations of individual LHCIIs between different conformational states, as has been observed by singlemolecule spectroscopy and has been suggested by MD simulations, do occur *in vivo*. The NMR experiments on whole thylakoid membranes containing LHCII are the first attempt, to the best of our knowledge, to obtain dynamic information of LHCII in a native thylakoid environment and under non-crystallizing conditions. Our NMR data results show that in a native setting where the proteins are embedded inside thylakoidal membranes, fast large-amplitude protein and pigment fluctuations are suppressed and flexible sites, involved in stabilization of the carotenoids, are motion-limited. While this suggests that spontaneous conformational fluctuations of LHCII are unlikely to occur in native membranes, differential NMR signals compared to those of liposome-embedded LHCII suggest that LHCII can be stabilized in different conformational states depending on its micro-environment. The thylakoid membrane is not a static architecture but in living organisms actively responds to changes in the light conditions. Thylakoid remodeling under light stress involves phosphorylation, membrane unstacking, super-complex reorganization and xanthophyll exchange, all of which may modify the protein landscape of LHCII and (temporarily) reduce the energetic transition barriers between distinct protein states. This work shows that by MAS NMR, structural dynamics can be uncovered from proteins, pigment and associated lipid components inside native, heterogeneous organelle membranes. The approach may open ways to explore how thylakoid plasticity controls the conformational states of the most abundant light-harvesting proteins, which are among the key players in photo regulatory processes.

## Supporting information

SI section

## ASSOCIATED CONTENT

Supporting Information. Fluorescence lifetime analysis and 77K fluorescence spectrum of LHCII proteoliposomes, chemical shift assignment tables and additional NMR spectra are provided in the Supporting Information section. The Supporting Information is available free of charge on the ACS Publications website.

## AUTHOR CONTRIBUTION

A.P. and F.A. designed the research, with the input of T.M. and M.B. F.A. and M.W. performed the experiments. G.P. and D.S. prepared the *U*-^13^C-LHCII complexes and F.A. performed the protein membrane reconstitution and prepared the *U*-^13^C-thylakoid membranes. F.A. and A.P. wrote the manuscript with input from all authors.

## ACKNOWLEDGEMENTS

We thank Emanuela Crisafi for assistance with the liposome preparations and Dr. Lijin Tian for assistance with the fluorescence experiments and the Biophysics Department of the VU University Amsterdam for use of their fluorescence spectrometry equipment. AP and FA were financially supported by a CW-VIDI grant of the Netherlands Organization of Scientific Research (NWO) under grant nr. 723.012.103. MEW is a recipient of a Natural Sciences and Engineering Research Council of Canada Postdoctoral Fellowship. This work was supported in part by uNMR-NL, an NWO-funded National Roadmap Large-Scale Facility of the Netherlands (grant number: 184.032.207).

