## Supplementary material for "Conformational dynamics of Light-Harvesting Complex II in a native membrane environment": SI section

Fatemeh Azadi-Chegeni^1^; Meaghan Ward^2^; Giorgio Perin^3†^; Diana Simionato^3^;Tomas Morosinotto^3^ ; Marc Baldus^2^; Anjali Pandit^1,*^

^1^ Leiden Institute of Chemistry, Dept. of Solid-State NMR, Leiden University, Einsteinweg 55, 2333CC, Leiden, The Netherlands

^2^ NMR Spectroscopy, Bijvoet Center for Biomolecular Research, Utrecht University, Padualaan 8, 3584 CH, Utrecht, The Netherlands

^3^ PAR-Lab (Padua Algae Research Laboratory), Dept. of Biology, University of Padova, Via Ugo Bassi 58B, 35121 Padova, Italy

†Current address: Dept. of Life Sciences, Imperial College London, Imperial College Rd., SW72BB, London, UK

**Table S1**

| Sample | A1 | τ_1_ (ns) | A2 | τ_2_ (ns) | A3 | τ_3_ (ns) | τ_av_ (ns) |
| --- | --- | --- | --- | --- | --- | --- | --- |
| ^13^C-^15^N LHCII in  α-DM | 62% | 3.8 | 18% | 1.5 | 20% | 0.1 | 3.5 |
| ^13^C-^15^N LHCII proteoliposomes | 1% | 7.2 | 65% | 0.7 | 34% | 0.3 | 0.7 |

Fluorescence lifetime analysis of *U*-^13^C-^15^N *Cr* LHCII in α-DM and in proteoliposomes.

**Table S2**

Assignment of mobile amino-acid residue types detected in the *J*-based INEPT-TOBSY spectrum of LHCII.

| Residue type | Cα [ppm] | Cβ [ppm] | Cγ [ppm] | Cδ [ppm] | Cε__[ppm] |
| --- | --- | --- | --- | --- | --- |
| Ala | 52.4 | 19.3 |  |  |  |
|  | 53.3 | 18.9 |  |  |  |
|  | 53.8 | 19.8 |  |  |  |
| Thr | 61.9 | 69.7 | 21.5 |  |  |
|  | 63.2 | 68.6 | 22.2 |  |  |
|  |  | 71.1 | 22.8 |  |  |
| Ser | 58.2 | 63.8 |  |  |  |
|  | 59.3 | 62.9 |  |  |  |
|  | 60.0 | 64.7 |  |  |  |
| Phe | 58.8 | 39.1 | 137.5 | 131.9 |  |
| Ile | 62.4 | 38.6 | 17.4 | 13.3 |  |
| Lys | 57.2 | 32.7 | 24.2 |  | 41.4 |
| Val | 63.2 | 31.9 | 20.6 | 19.3 |  |
| Glu | 57.3 | 29.7 |  |  |  |
| Asn | 53.1 | 38.9 |  |  |  |
| Leu | 56.1 | 42.5 |  |  |  |
| Pro | 63.8 | 31.5 | 26.4 | 48.9 |  |
|  |  |  | 27.4 | 50.8 |  |
|  |  |  | 26.9 | 49.5 |  |
| Asp/Leu | 54.2 | 41.0 |  |  |  |

**Table S3**

Assignment of Chlorophyll tail ^13^C resonance signals detected in the *J*-based INEPT-TOBSY spectrum of LHCII.

| Carbon atom | Chemical shift (ppm)  Chl i [ppm] Chl ii[ppm] |
| --- | --- |
| P1 | 60.8 62.9 |
| P2 | 121.6 126.1 |
| P3 | 140.8 143.5 |
| P3a | 18.4 18.4 |
| P4 | 42.5 42.1 |

**Table S4**

Assignment of ^13^C lipid resonance signals detected in the *J*-based INEPT-TOBSY spectrum of LHCII and in the spectrum of thylakoid membranes.

| Carbon atom | Chemical shift (ppm)  MGDG | DGDG |
| --- | --- | --- |
| C1 | 105.9 | 101 |
| C2 | 75.4 | 74.3 |
| C3 | 73.4 | - |
| C4 | 68.9 | 77.7 |
| C5 | - | 71.1 |
| C6 | 65 | 54.6 |
| C7 | - | - |
| C8 | 130.2 | 130.2 |
| C9 | 132.1 | 132.1 |
| C10 | 29.6 | 29.6 |
| C11 | 27.9 | 27.9 |
| C12 | 130.7 | 130.7 |
| C13 | - | - |
| C14 | 27.8 | 27.8 |
| C15 | 129.6 | 129.6 |
| C16 | 133.8 | 133.8 |
| C17 | 23.1 | 23.1 |
| C18 | 16.66 | 16.66 |

**Figure S1**

LHCII structure and abundancy in thylakoid membranes. A: Protein structure. B: LHCII structure including the pigments, with Chl *a* in green, Chl *b* in blue, lutein in red and neoxanthin in purple and violaxanthin in yellow. C: top view of trimeric LHCII. D: Sucrose gradient of *Cr* thylakoid membranes after solubilizing with 0.6% α-DM, showing the fraction of trimeric LHCII. E: SDS page gel image of the *Cr* thylakoid membranes and of isolated LHCII.

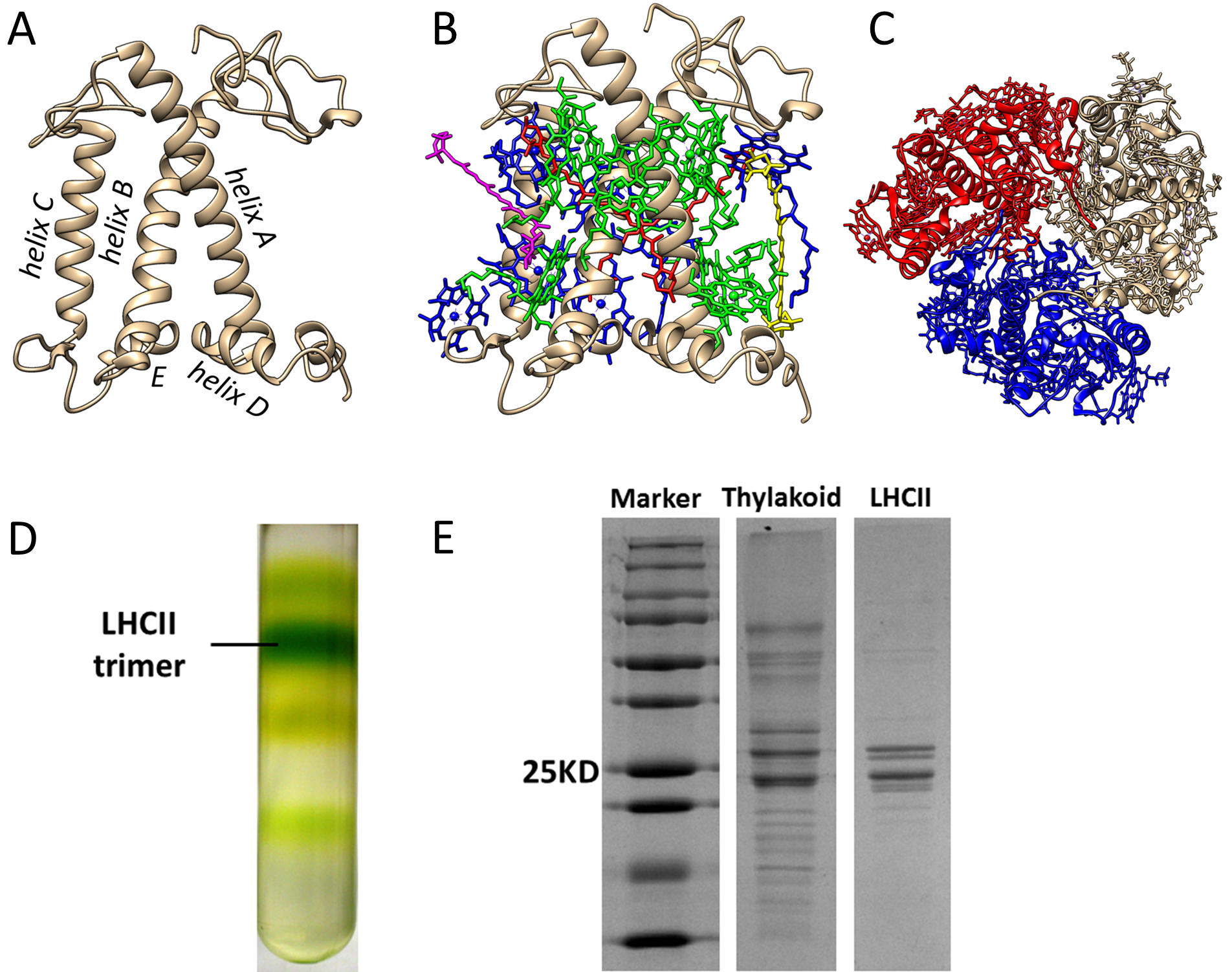

**Figure S2**

A: Example of thylakoid extraction using a layered sucrose gradient. a. eye spot containing β- carotenes; b. thylakoid membranes; c. cell walls and unbroken cell material. Band b was extracted with a syringe and contained the purified thylakoid fraction.

B: Absorption spectra of ^13^C *Cr* thylakoid membranes. The Q_y_ absorbance bands of Chl*a* and *b* at are distinguished at 672 and 650 nm respectively, and carotenoids and Chl higher-energy states contribute to the spectrum in the region between 400 and 500 nm.

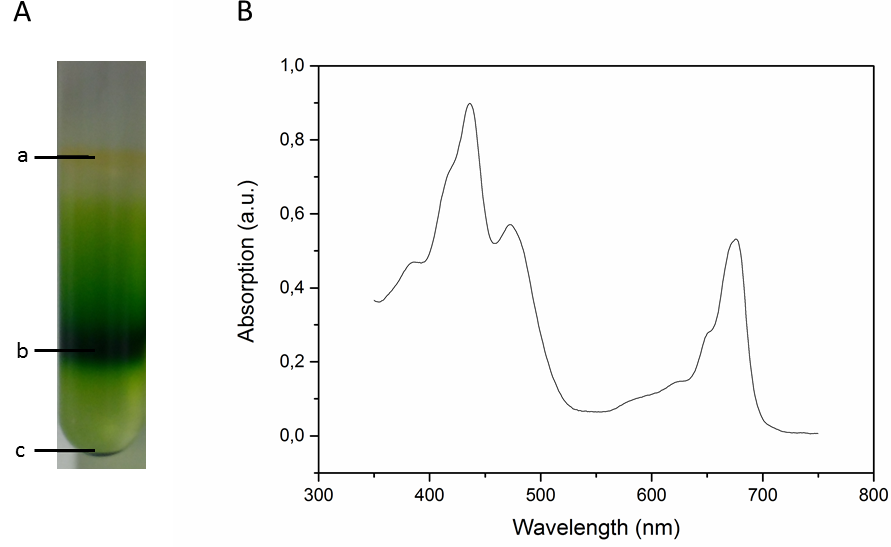

**Figure S3**

77K fluorescence spectrum of LHCII in α-DM (black) and of LHCII proteoliposomes (red). The band at 700 nm is a signature of LHCII aggregation in the proteoliposomes.

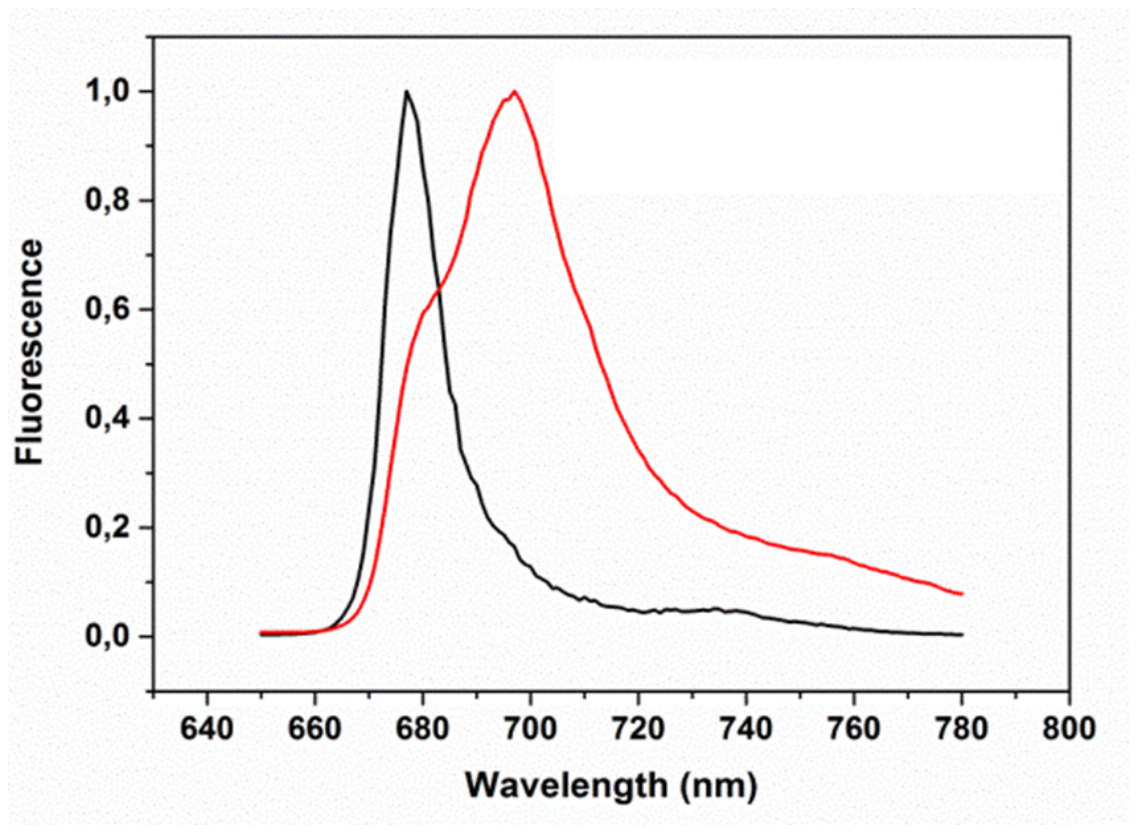

**Figure S4**

Time-resolved fluorescence decay of LHCII in α-DM (green) and of LHCII proteoliposomes (black). Dotted lines in blue and red are exponential fit curves using three components.

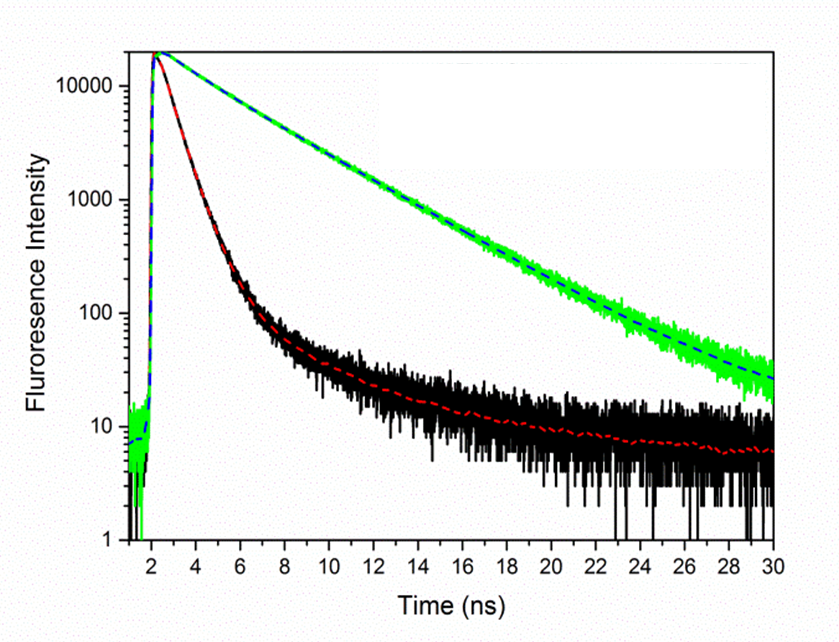

**Figure S5**

CP-PARIS ^13^C-^13^C spectrum of LHCII proteoliposomes, aliphatic region. The spectrum was collected with a mixing time of 30 ms at 17 kHz MAS at a set temperature of -18 ^o^C.

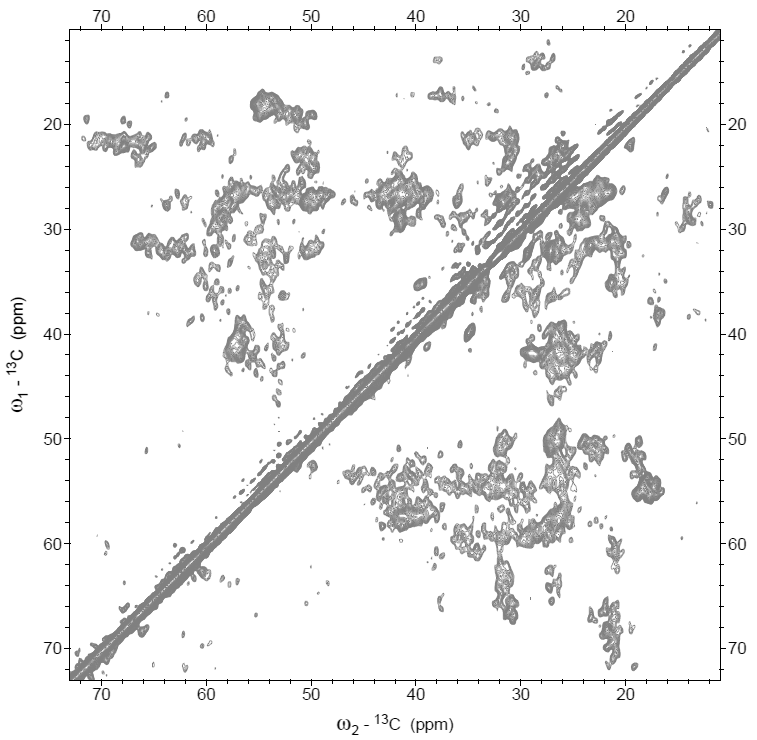

**Figure S6**

Structure of Lhcbm1, highlighting Ala (A), Gly (B), Thr (C) and Ser (D) residues.

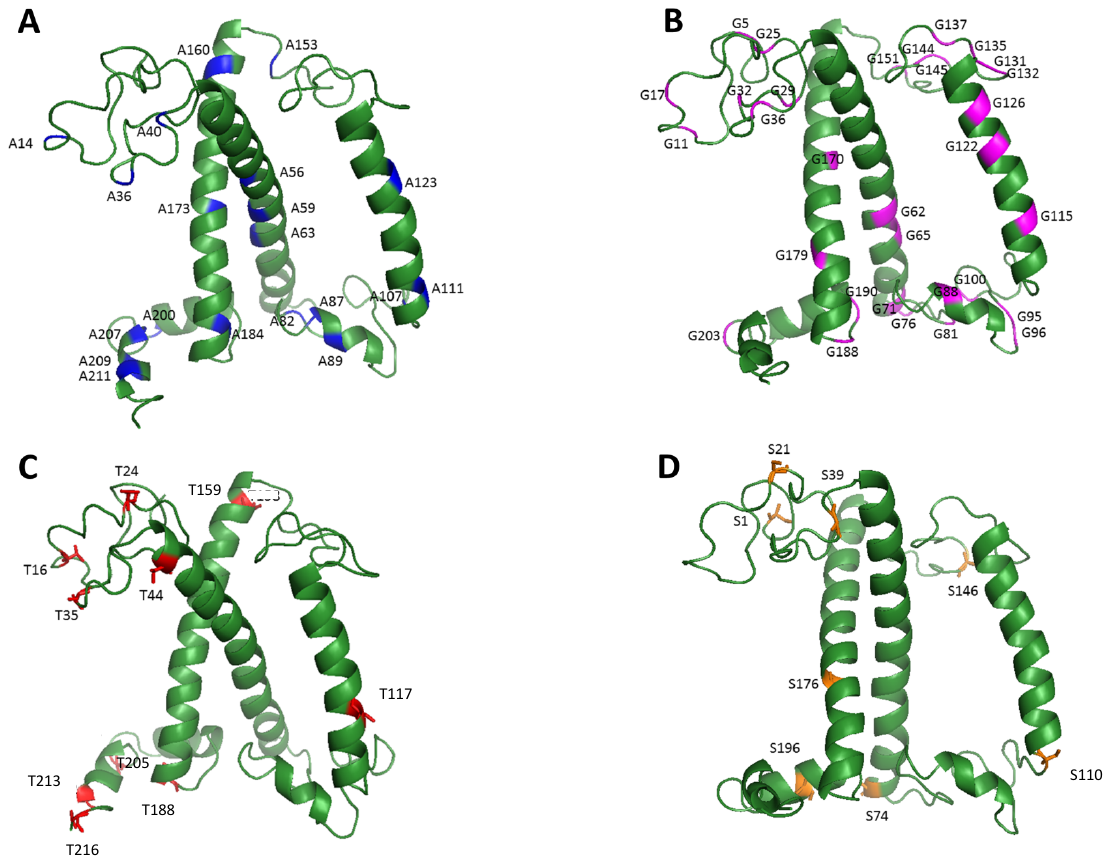

**Figure S7**

Helix and coil percentages according to the NMR spectrum, *Cr* LHCII homology model based on Lhcbm1 (model1), Lhcbm2/7 (model2) or Lhcbm3 (model3) and according to the full amino acid sequence of Lhcbm3 (sequence3). The Lhcbm1 model lacks the N-terminal amino acids that are present in the sequence. The structural model for Lhcbm2 contains the full protein sequence as this polypeptide has no N tail.

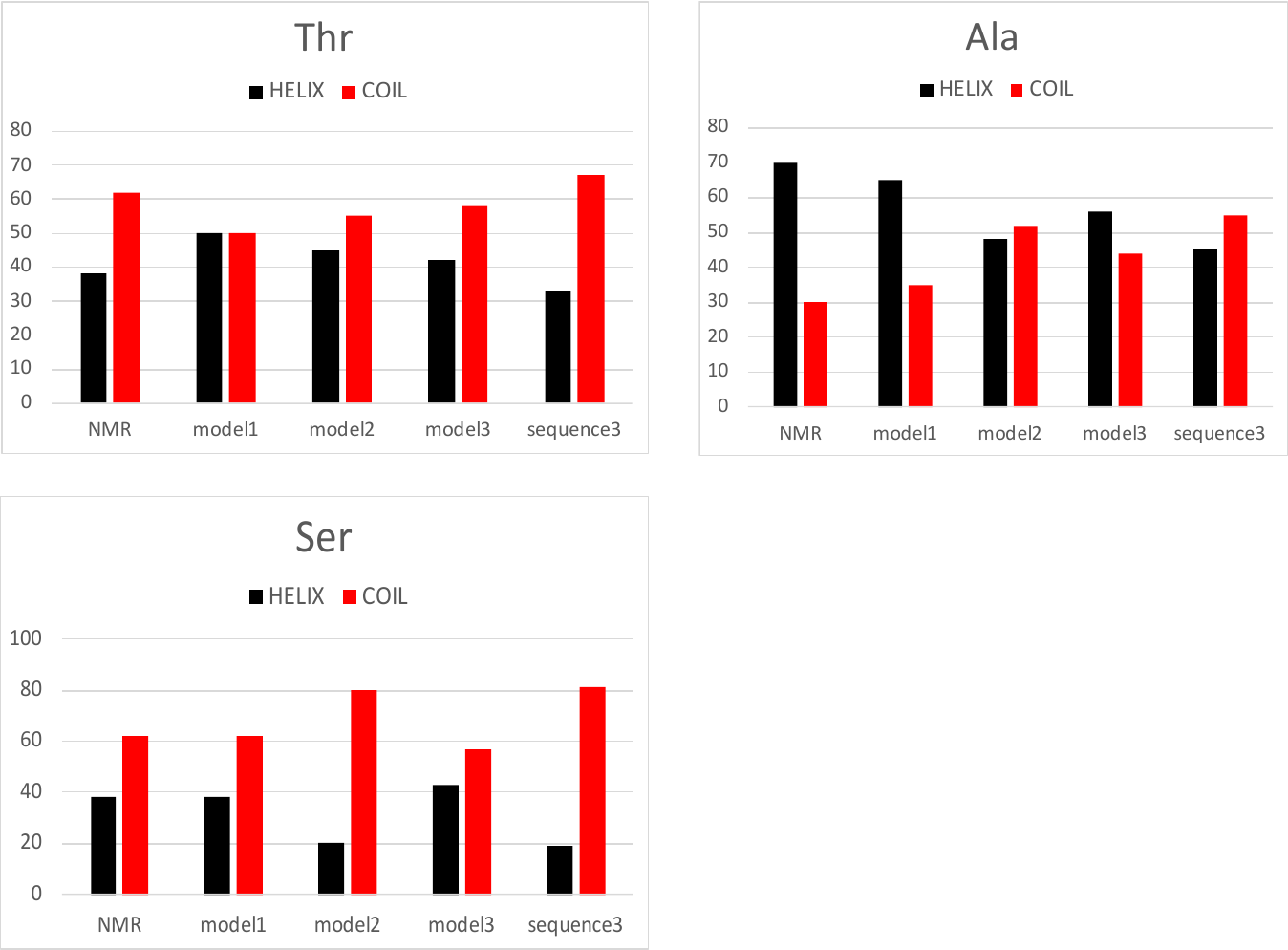

**Figure S8**

Comparison of 1D-^13^C CP-MAS and direct excitation (DP)-MAS spectra. Overlaid ^13^C CP (black) and direct excitation (DP, green) spectra of thylakoid membranes containing LHCII (A) and of LHCII proteoliposomes (B). The spectra were collected with 512 scans, 2 second of recycle delay, 80 ms acquisition time at 14 kHz. For CP-MAS experiments, the mixing time was set to 1 millisecond. The set temperature was -3 ^o^C.

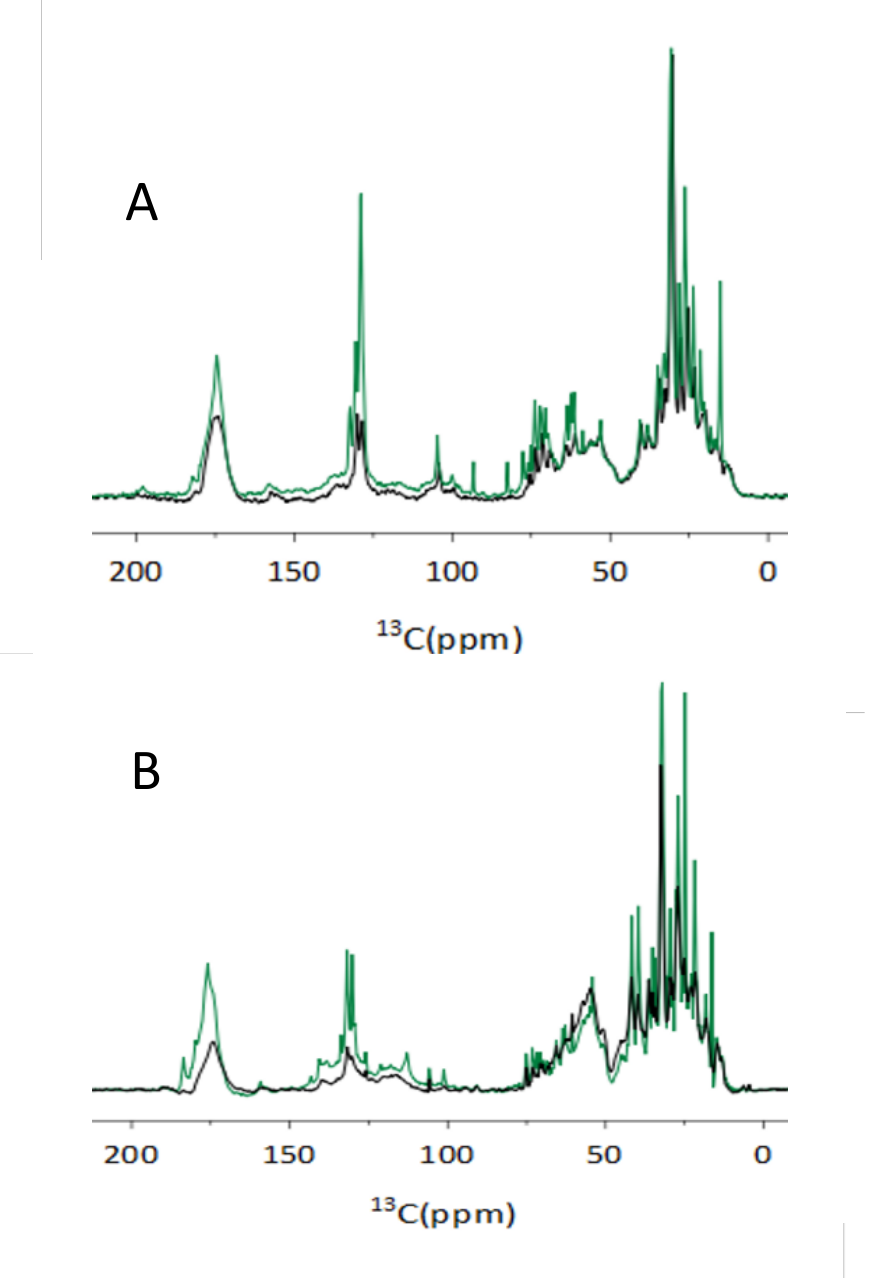

**Figure S9**

^13^C-^13^C CP-PARIS spectrum of LHCII proteoliposomes (red) and of thylakoid membranes containing LHCII (black). Spectra were collected with a mixing time of 30 ms at 17 kHz MAS at a set temperature of -18 ^o^C.

Below: Aromatic region showing correlation signals of Chls and carotenoids (Car).

Next page: Selective 1D slices of the ^13^C-^13^C CP-PARIS spectrum of LHCII proteoliposomes (red) and of thylakoid membranes containing LHCII (black) with Ala, Ser and Thr peaks in helix and coil regions.

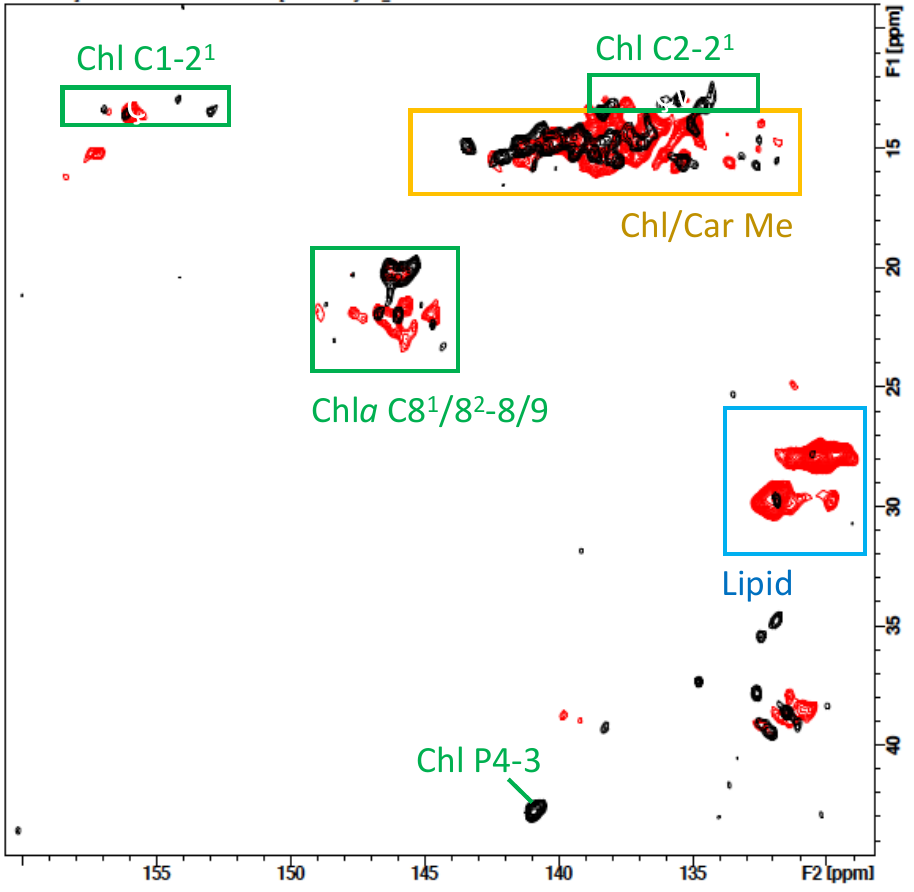

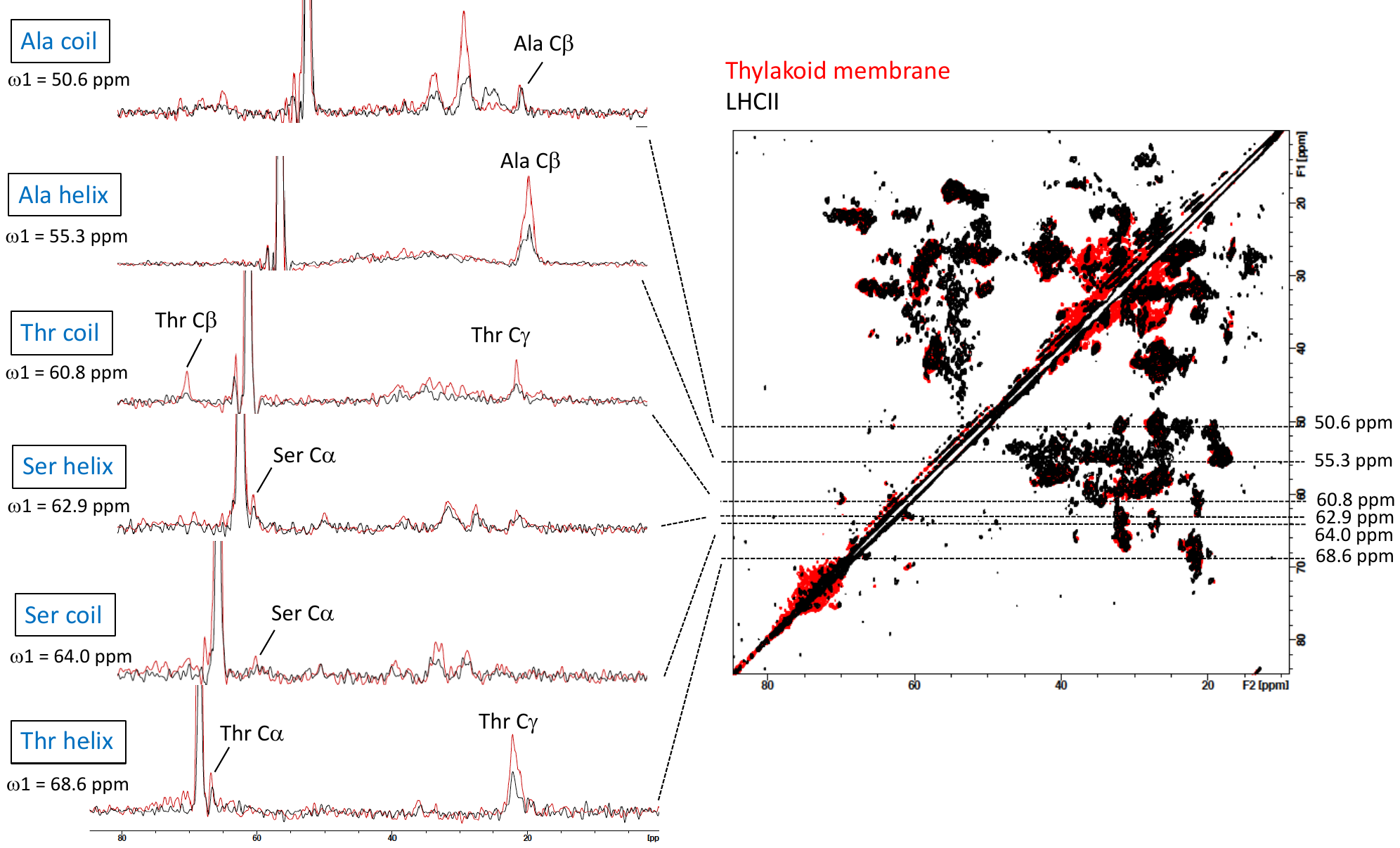

**Figure S10**

CP-PARIS ^13^C-^13^C spectrum of LHCII proteoliposomes (black) overlaid with SHIFTX2-generated protein correlations prediction of Lhcbm1 (cyan) and Lhcbm2 (orange). The spectrum was collected with a mixing time of 30 ms at 17 kHz MAS at a set temperature of -18 ^o^C.

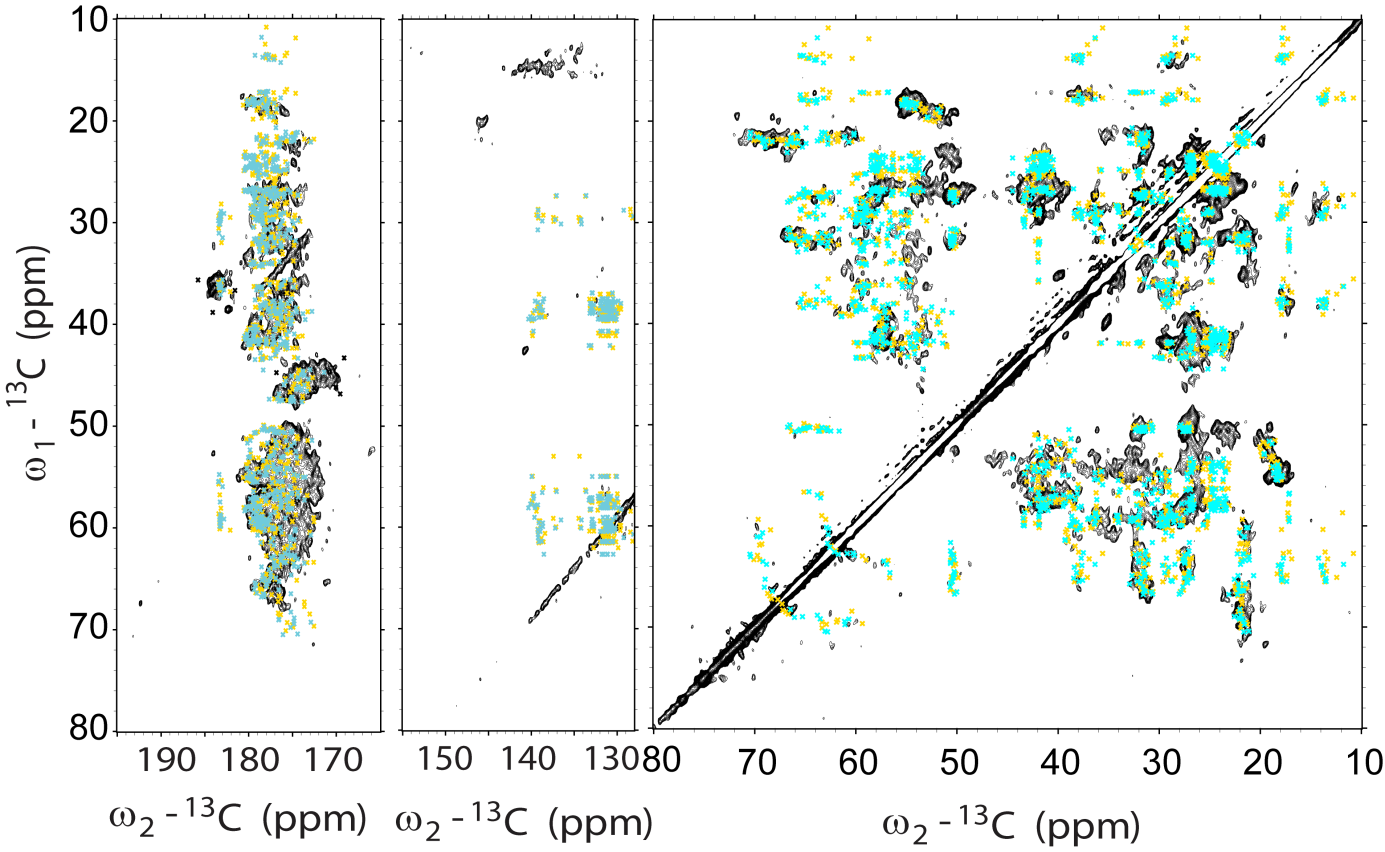

**Figure S11**

NCA ^15^N-^13^C spectrum of LHCII proteoliposomes (black) overlaid with predicted correlations of Lhcbm1 (red dots) and Lhcbm2 (green dots). The NCA spectrum was collected at 14 kHz MAS frequency at -18 ^o^C, using a mixing time of 800 μs ^1^H-^15^N CP step, 4.2 ms for ^15^N-^13^C and the number of scans was set to 704. Significant deviations between predicted and experimentally observed cross peaks are observed for Val106, Ile111, Thr188 and Thr213 in Lhcbm1, and in Gly20, Ile114 and Thr216 in Lhcbm2.

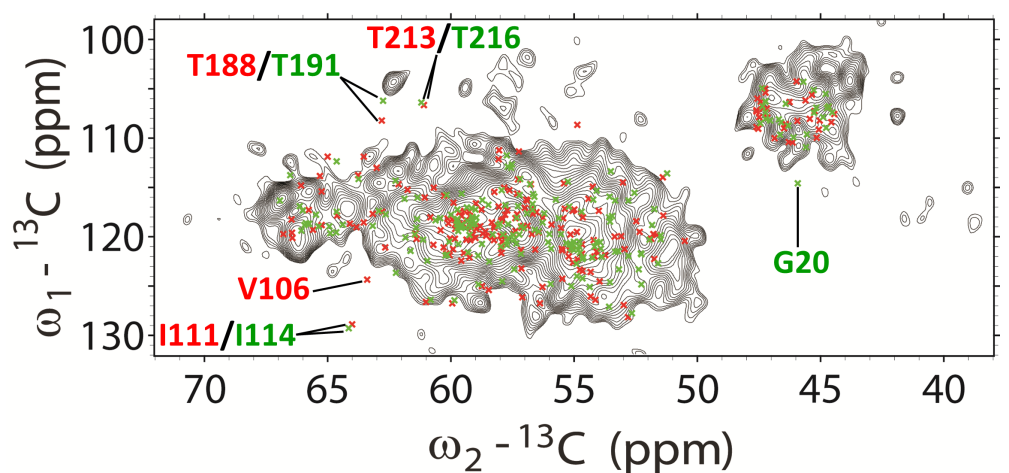

**Figure S12**

A-D: Overlay of the INEPT-TOBSY spectrum of LHCII proteoliposomes (black) and of thylakoid membranes containing LHCII (red). Protein assignments are indicated in black, Chl assignments in green and lipid assignments in blue. Spectra were collected at -3 ^o^C. E: Chemical structure of MGDG, highlighting the lipid atom types that could be distinguished in the INEPT-TOBSY spectrum.

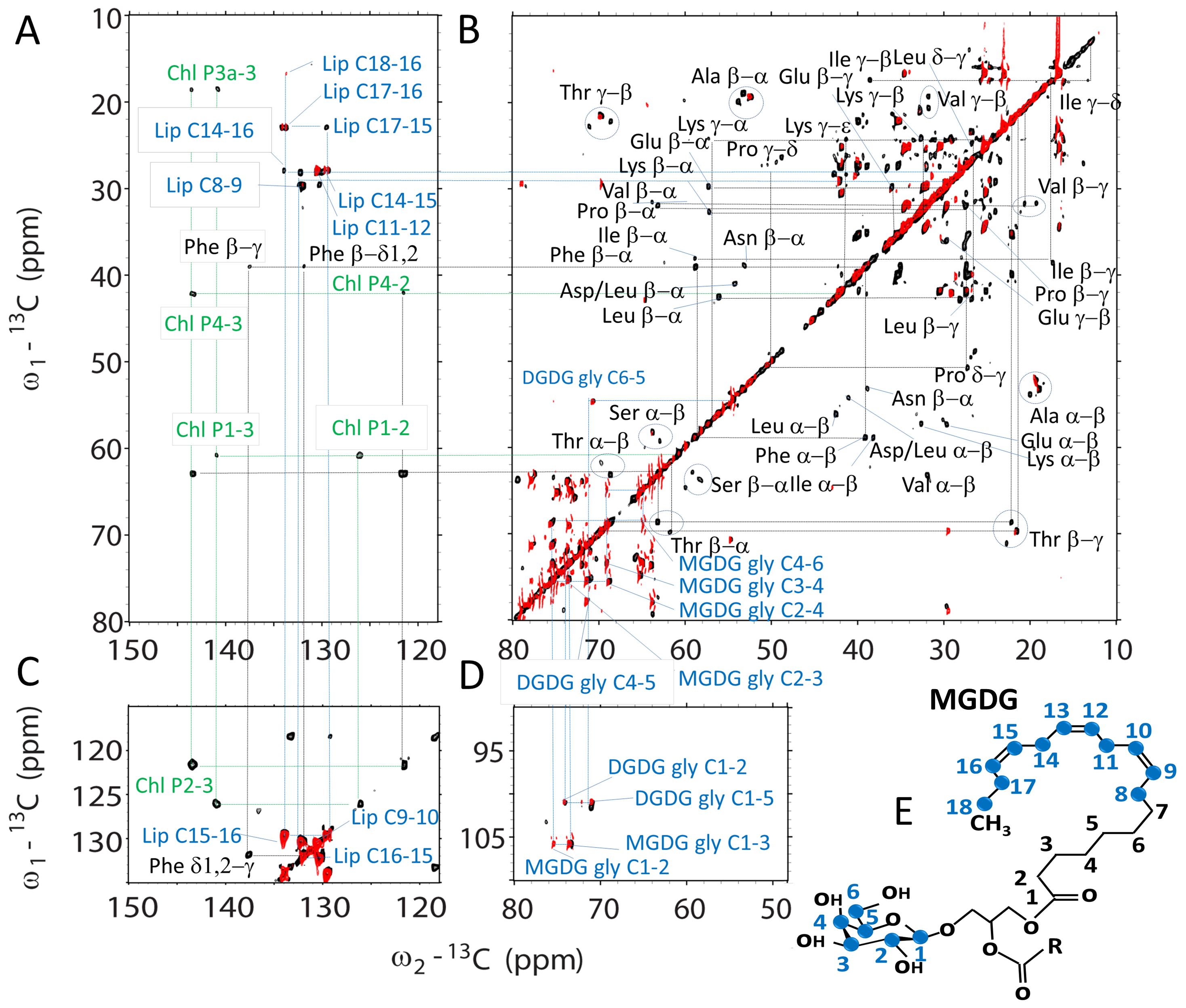

**Figure S13**

Overlaid ^13^C-^13^C PARIS (black) and ^13^C-^13^C INEPT-TOBSY (blue) spectra of LHCII proteoliposomes. The insets show the Ala, Ser and Thr signals. The CP-PARIS spectrum was recorded at a set temperature of 255 K and the INEPT-TOBSY spectrum at a set temperature of 270 K. One Ser and two Ala peaks overlap in the two spectra.

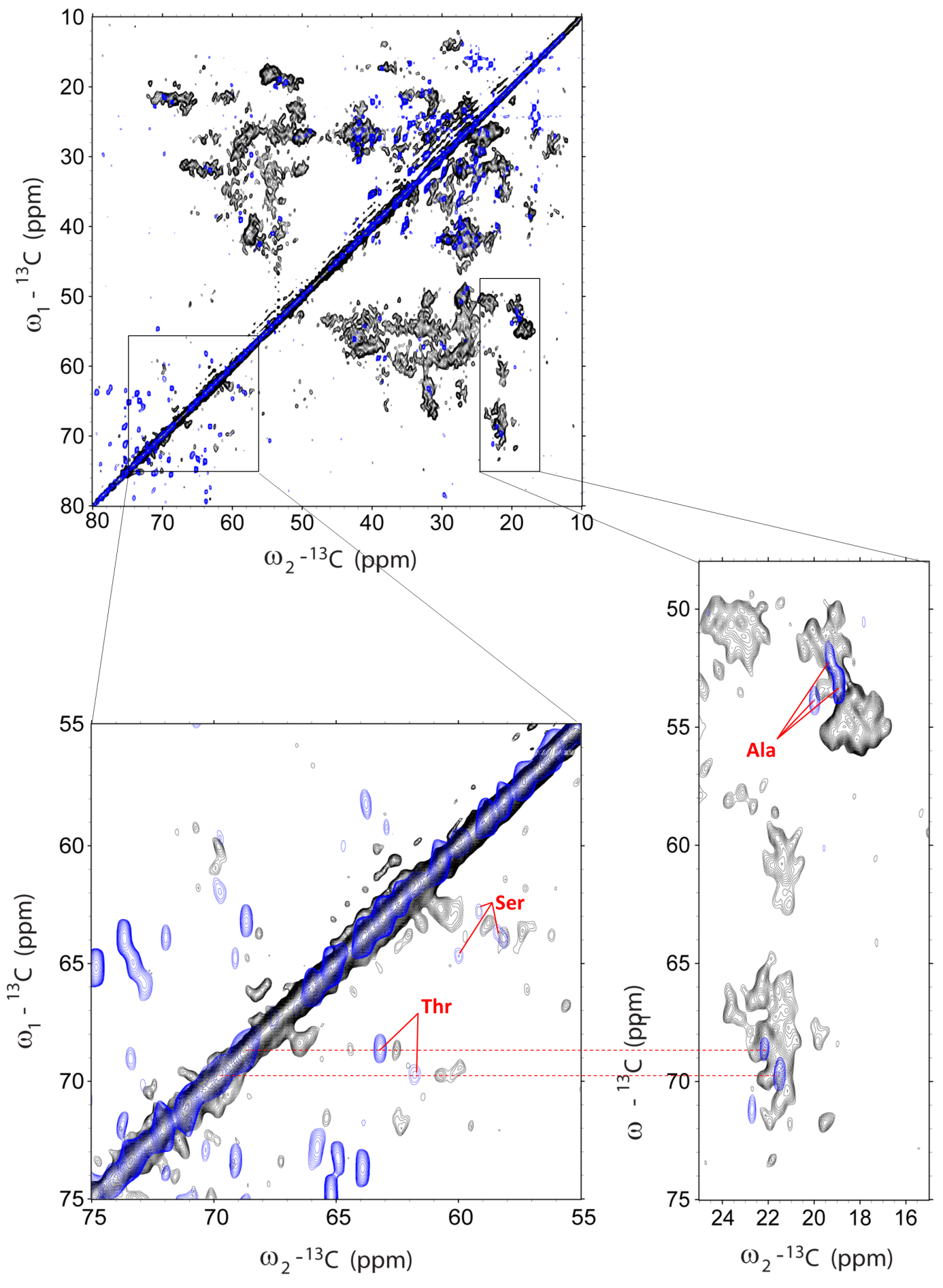

**Figure S14**

Overlaid INEPT-TOBSY (red) and DP-PARIS (blue) spectrum showing correlations of Chl phytol (P) carbon atoms. The spectra were collected at -3 ^o^C and 14 kHz MAS. For the TOBSY spectrum, a mixing time of 6 ms was applied.

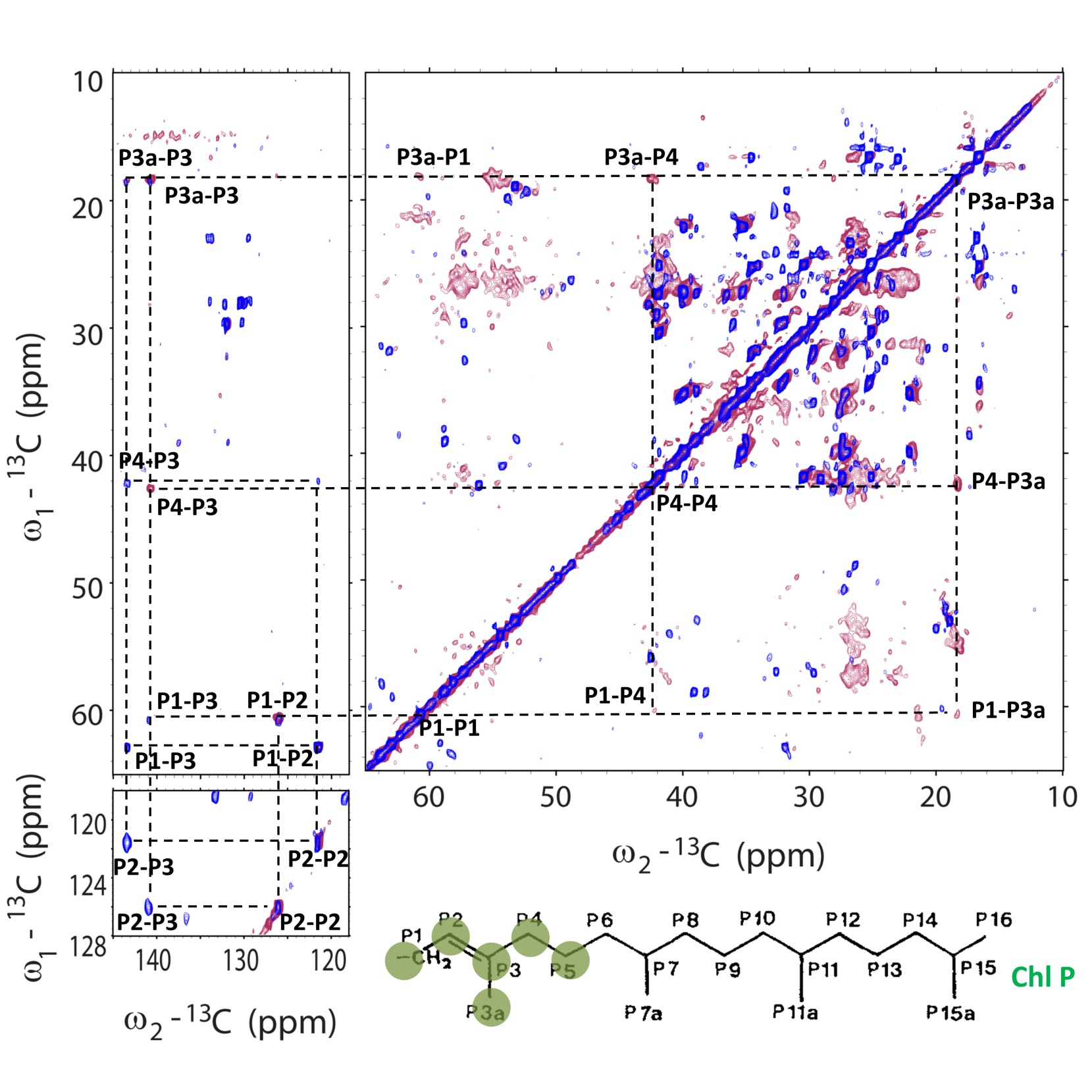

**Figure S15**

NMR comparison of structure-predicted and experimental NMR chemical shift correlations. A-D: ^13^C-^13^C CP-PARIS spectrum of thylakoid membranes containing LHCII (red) with the LHCII proteoliposome spectrum (black) overlaid. The insets show Ala (B), Thr (C) and Ser (D) spectral regions overlaid with chemical-shift predictions of Lhcbm1 (cyan crosses) and Lhcbm2 (black crosses). Predicted shifts that significantly deviate from experimental correlations are highlighted in yellow.

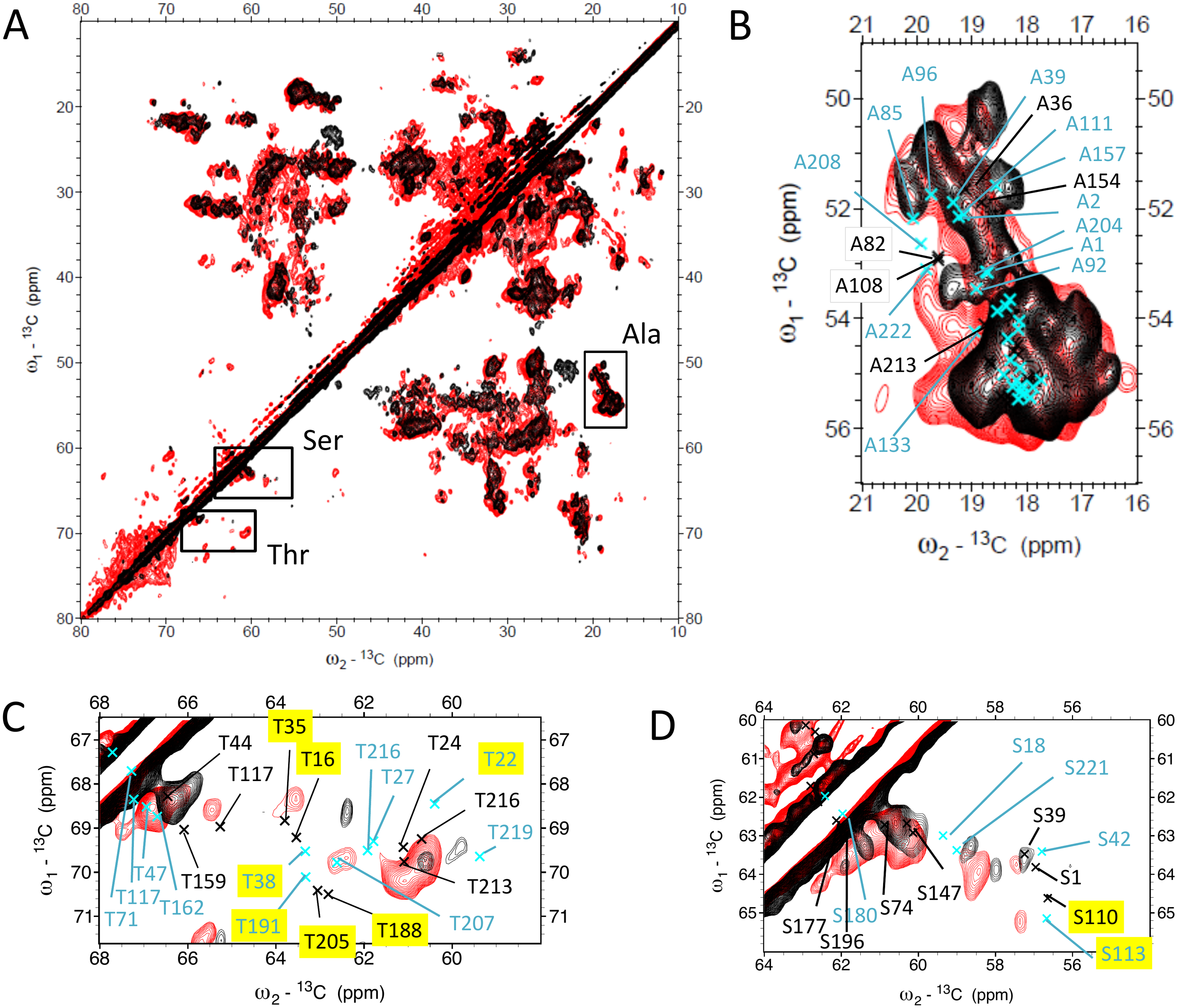
